# Growth history leaves a geometric trace in puzzle cells

**DOI:** 10.1101/2023.10.01.560343

**Authors:** Nicola Trozzi, Brendan Lane, Alice Perruchoud, Frances Clark, Lukas Hoermayer, Andrea Meraviglia, Tammo Reichgelt, Adrienne H.K. Roeder, Dorota Kwiatkowska, Adam Runions, Richard S. Smith, Mateusz Majda

## Abstract

Puzzle-shaped epidermal cells not only reduce mechanical stress during organ growth but also record the expansion history of the tissue in their outlines. By combining mechanical simulations with time-lapse imaging, we show that transitions from directional to isotropic expansion induce new lobes to form along the previous growth axis, and that switching the order of anisotropic and isotropic phases yields hybrid shapes that reliably preserve those transitions. In maize, model predictions and live imaging coincide precisely, and in *Arabidopsis*, final lobe patterns correlate more with growth history than with cell size alone. Genetic or pharmacological disruption of lobe formation constrains leaf expansion or drives compensatory elongation, which underscores a mechanical function. A broad survey of living and fossil vascular plants reveals that the mechanism to make puzzle-shaped cells is both widespread and developmentally plastic, suggesting that single snapshots of leaves can give insight into their growth history. Together, these findings demonstrate that puzzle cells transform cell geometry into a living record of how tissues grow.

## Introduction

The shapes of plant organs, whether broad or elongated, are often reflected in the shapes of individual cells, suggesting a close relationship between organ growth patterns and cellular morphology (Hülskamp et al., 1998; Zuch et al., 2022; Hu et al., 2024). Studies using *Arabidopsis thaliana* mutants have demonstrated that alterations in organ shape are accompanied by corresponding changes in cell shape and vice versa, highlighting the interdependence of growth dynamics at multiple scales (Qiu et al., 2002; Hake et al., 2004; Vlad et al., 2014; Tasker-Brown et al., 2024). In the case of simple-shaped cells, the relationship is intuitive; elongated organs tend to have long, thin cells. However, when epidermal tissues make puz-zle-shaped cells, this relationship appears at first glance to be lost. Integrating experimental biology with theoretical modeling has advanced our understanding of how genes, mechanics, and growth interact. This approach emphasizes the need to view tissues and organs as integrated systems rather than isolated components. Although the molecular regulation of growth occurs at the cellular level, the specified growth of individual cells must conform to their neighbors, and the resultant growth that emerges is often different due to mechanical conflicts (Rebocho et al., 2017). This idea also applies to cell shape, which emerges from collective tissue dynamics and not solely from the molecular regulation of individual cells (Sapala et al., 2018; Vőfély et al., 2019; Majda et al., 2021).

Endoreduplication, characterized by successive rounds of DNA replication without cell division, results in significant cell enlargement, increasing the mechanical demands on cell walls to withstand the elevated stress from turgor pressure in larger cells (Roeder et al., 2010; Edgar et al., 2014). In stems and hypocotyls, cells typically elongate along the main growth axis (Gendreau et al., 1997; Baskin, 2005; Cosgrove, 2022). This elongated shape naturally limits the stress, which can be approximated by the Largest Empty Circle (LEC), the largest circular area within a cell without internal anticlinal walls. Although the LEC is computed on 2D outlines, prior 3D finite element modeling has shown that it captures the maximal unsupported span of the outer periclinal wall under turgor, integrating curvature and depth effects (Sapala et al., 2018). At the stage when puzzle cells emerge, curvature differences between the abaxial and adaxial surfaces are minor compared with the strong local curvature of individual cells, so the LEC provides a robust approximation of the stresses experienced by pavement cells. However, nearly isotropic growth in broad aerial organs like leaves presents a distinct challenge, as cells expanding isotropically risk developing large, mechanically vulnerable areas that could rupture due to internal stress (Hamant and Traas, 2010; Kierzkowski and Routier-Kierzkowska, 2019; Malivert et al., 2021). To address this, epidermal cells frequently adopt a puzzle-shaped morphology with interlocking lobes. These lobes decrease the LEC, reduce the mechanical stress, and help maintain tissue integrity under tension (Majda et al., 2017; Sapala et al., 2018; Bidhendi et al., 2019; Bidhendi and Geitmann, 2019; Bidhendi et al., 2023). Finite element modeling and 3D stress analysis support a functional role for these unusual shapes, showing that lobed cells experience reduced wall stress during isotropic expansion. This suggests that puzzle shape formation reflects a mechanical adaptation to accommodate endoreduplication-induced cell enlargement and organ scale growth dynamics, positioning mechanical stress as a key constraint in shaping epidermal cell morphology (Sapala et al., 2018).

At the cellular level, plant cells are tightly connected by their cell walls, meaning that their growth must be coordinated to maintain tissue integrity. This coordination is influenced by the interplay between internal turgor pressure and the mechanical properties of the cell wall, which together constrain how neighboring cells can grow relative to each other. Turgor pressure generates tensile in-plane stress across the cell wall, which resists expansion while the wall is remodeled in a controlled fashion. This remodeling includes the regulated sliding of cellulose fibers at specific attachment points, allowing the cell to grow without compromising its structure (Zhang et al., 2021; Cosgrove, 2022; Coen and Cosgrove, 2023). Cortical microtubules guide the localized deposition of cellulose microfibrils, thereby stiffening specific regions of the cell wall and reinforcing cell structure. In puzzle-shaped cells, microtubules accumulate in narrow neck regions between protruding lobes, restricting local expansion and shaping the lobed contour, whereas in elongated cells they align transversely to limit lateral growth (Baskin, 2005; Panteris and Galatis, 2005; Paredez et al., 2006; Sampathkumar et al., 2014). One hypothesis tested in a spring-based model was that localized growth restrictions create indentations that then attract microtubules and become reinforced, establishing a feed-back loop. Due to growth conflicts, neighbor cells will have lobes that correspond to the indentations, with coordination emerging as a mechanical effect (Sapala et al., 2018). Computational simulations based on these ideas have successfully reproduced a variety of cell shapes and the significant variation found in nature, demonstrating their effectiveness in modeling the dynamics of puzzle cell shape acquisition. Although unintuitive, the model shows that anisotropic growth leads to long, thin cells without lobes, whereas nearly isotropic growth yields more complex puzzle-shaped cells.

If puzzle cell shape is an adaptation to a mechanical constraint based on stress, then one prediction would be that most plants would possess the ability to make puzzle cell shapes. A positive correlation between cell area and lobe number has been observed in *Arabidopsis* and several other species (Sapala et al., 2018). However, in a broader survey of vascular plants (Vőfély et al., 2019), no correlation between cell area and cell lobing was found, and the average lobeyness remained low even in species with large cells, indicating that highly lobed cells, like those in *Arabidopsis* leaves or cotyledons, are relatively rare. They proposed that epidermal cell shapes do not have a conserved function across vascular plants, but rather may serve diverse functional roles across taxa, representing species-specific adaptive strategies shaped by different ecological and evolutionary pressures.

In this work, we investigate whether the ability to form puzzle-shaped cells is broadly conserved across vascular plants, and how developmental stage, organ identity, and environmental conditions influence their occurrence. By combining modeling, live imaging, and a large-scale comparative survey, we aim to reveal how growth history and mechanical constraints interact to shape epidermal cell morphology.

## Results

### Impact of organ growth dynamics on puzzle cell formation

The spring model proposed by Sapala et al. (2018) provides a mechanistic explanation for the formation of shapes that maintain small LECs, ranging from puzzle-shaped to elongated cells. This model offers a framework not only for understanding how shape emerges at the cellular level but also provides an unexplored opportunity to study the interaction of mechanical constraints and growth dynamics at the organ scale. To empirically validate this model, we ex-amined the distribution of cortical microtubule orientation in experimentally observed epi-dermal pavement cells. By analyzing the cell wall morphology and microtubule patterns (**Figure 1A**) and using these observations to create segmented cell models for simulation, we successfully replicated the microtubule dynamics predicted by the model. Simulated connec-tions qualitatively correspond to cortical microtubules and downstream cellulose deposition were more concentrated in the indentations and notably sparse in the lobed regions of the cells (**Figure 1A, B**). These results demonstrate that the Sapala et al. (2018) model can gen-erate realistic puzzle cell shapes with microtubule patterns that match experimental observa-tion.

**Figure 1.**
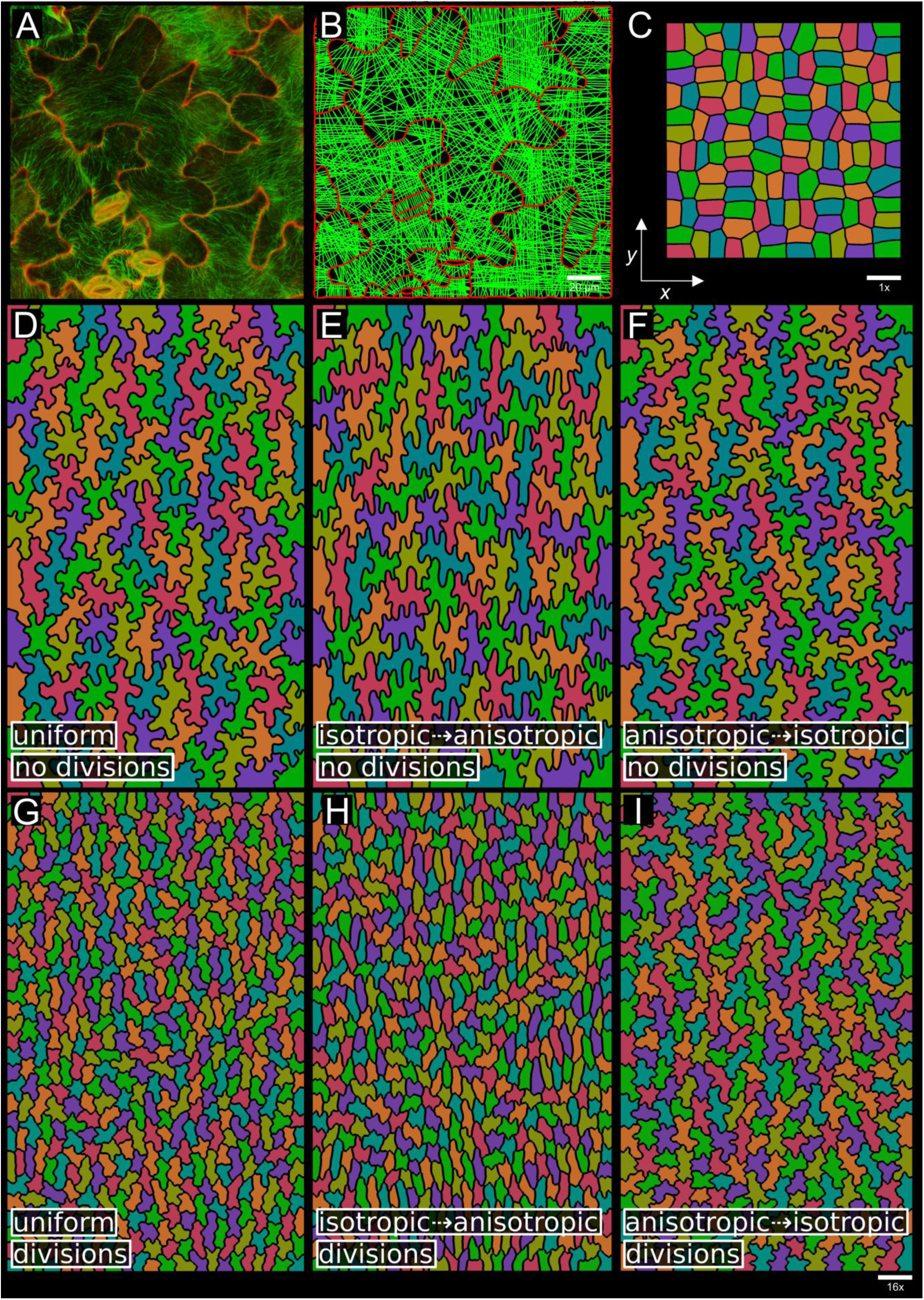
Growth distribution over time influences the shapes of pavement cells. (**A**) Confocal image representing epidermal pavement cells from a fully developed 3-week-old leaf of *Arabidopsis thaliana*. The cell wall was visualized using propidium iodide staining (red), and the microtubules were visualized using fluorescent TUA6-CFP (green). (**B**) A visualization of the placement of connections across the cells for the cell shapes in **A** in the computational model. (**C-F**) Model simulation of puzzle cell formation across various growth fields. (**C**) The initial template. (**D-F**) Cell shapes evolved over 300 growth iterations to the same final template size, 16 times larger in y and 10 times larger in x than the initial template. (**D**) Uniform weakly anisotropic growth with puzzle-shaped cells. Growth rates in the x and y directions are different but constant over time. (**E**) Initially isotropic growth followed by a short period of anisotropic growth in the y direction only (5/6 vs 1/6 of the simulation time) resulting in puzzle-shaped cells whose lobes are largely oriented vertically. (**F**) The same simulation with the growth reversed, a short burst of anisotropic growth followed by more isotropic growth, resulting in puzzle-shaped cells with a subtle bias in the horizontal orientation. (**G**-**I**) same as (**D**), (**E**), and (**F**) respectively, with divisions implemented that cease after the first 20 of 100 growth steps. Scale bars, as indicated.

Growth anisotropy significantly influences epidermal cell shape, generally producing elon-gated cells under anisotropic growth and puzzle-shaped cells under isotropic growth (Sapala et al., 2018). Building upon this finding, we further explored how the sequence of anisotropic and isotropic growth phases affects cell morphology, even when the total cumulative growth is the same. We conducted computational simulations starting from an initial template of iso-diametric cell shapes (**Figure 1C**), expanding the cells 16 times vertically and 10 times hori-zontally (**Suppl Figure 1**). In the first scenario, we applied uniform growth rates across all cells with an anisotropic ratio of 1.6:1, resulting in slightly elongated cells with lobes similar to the isotropic case (**Figure 1D, Suppl. Movie 1**), closely resembling cells observed in the proximal and distal abaxial regions of the leaf in *Arabidopsis* (**Suppl. Figure 2**). In a second scenario, we initially imposed isotropic growth followed by a brief period of vertical aniso-tropic growth (approximately 5/6 isotropic and 1/6 anisotropic), which led to vertically ori-ented lobes (**Figure 1E, Suppl. Movie 2**). Conversely, reversing the growth sequence, start-ing with anisotropic growth followed by isotropic growth, resulted in horizontally biased lobes (**Figure 1F, Suppl. Movie 3**). These simulations indicate that differences in the tem-poral dynamics of growth anisotropy, even under conditions of identical cumulative growth, can plausibly lead to significant variations in cell shape. Furthermore, isotropic tissue growth tends to yield more complex, puzzle-like cell morphologies, while anisotropic growth pro-duces simpler, elongated forms. These observations suggest that final cell shape reflects the cumulative outcome of dynamically evolving growth patterns, with changes in local growth anisotropy influencing morphogenesis in real time. This provides insight into how organ and cell shape interact when cells do not have simple shapes. We next turned to the role of prolif-eration, asking whether continued or halted division alters final cell size and lobing patterns.

### The interaction of cell division and growth

If the purpose of puzzle-shaped cells is to mitigate stress as the cells become large, one strategy would be for the plant cells to keep dividing. Some species appear to do this, making simple polygonal cell shapes (Remmler and Rolland-Lagan, 2012; Sapala et al., 2018; Vőfély et al., 2019; Liu et al., 2021). In plants that do make puzzle cell shapes, different developmental contexts can make different cell shapes, even if the growth dynamics are the same. One example is the abaxial vs abaxial sides of the *Arabidopsis* leaf, which show differences in puzzle cell shape (**Suppl. Figure 2**) (Panteris and Galatis, 2005; Vőfély et al., 2019; Liu et al., 2021). To explore the impact of the timing of the cessation of cell division, we ran the models with different growth dynamics (**Figure 1D-F**) but allowed cell division for the initial 20 of the 100 simulation steps (**Figure 1G-I, Suppl Movie 4-6**). Although not a large proportion of the simulation time, it has a substantial effect on the final cell shape. As expected, more divisions result in smaller and overall, less lobey cells. Less intuitively, it also affects the bias in lobing, which in some cases increases. In the simulation with first isotropic then anisotropic growth, the lobes appear almost exclusively in the vertical direction (compare **Figure 1E** and **1H**). In the uniform case, the model approximates the differences in cell shape between the abaxial and adaxial sides of an *Arabidopsis* leaf (compare **Figure 1D, G** with **Suppl. Figure 2**). This suggests that the difference in cell lobing between the abaxial and adaxial sides of the leaf could be explained by the timing of the entry into the endoreduplication cycle.

Experimental manipulation of division timing supports the model’s prediction that increased proliferation reduces lobeyness. The LOSS OF GIANT CELLS FROM ORGANS (LGO) regulator of endoreduplication affects this timing (Schwarz and Roeder, 2016). In the *lgo-2* mutant, which undergoes less endoreduplication than Col-0, cells divided more frequently and remained small and less lobed (**Suppl. Figure 5A, B**). In contrast, LGO overexpression produced larger cells with pronounced lobes (**Suppl. Figure 5C**). These observations are consistent with the idea that extended proliferation limits the expansion phase, restricting lobe development, and that changes in lobeyness arise indirectly from altered growth dynamics downstream of endoreduplication (**Suppl. Figure 5D**). While these *Arabidopsis* models reveal how cell division timing influences lobing under different growth patterns, we next asked whether similar principles apply in a species with strong directional growth.

### Experimental validation of growth history

We focused on maize (*Zea mays*), whose elongated leaves contain pavement cells with lobes oriented mainly in the transverse direction, an arrangement that is unexpected given the leaf shape, which suggests growth anisotropy should favor the longitudinal axis. To determine the growth rates and directions that produce these cell shapes, we used time-lapse experiments that obtained sequential replicas of cell patches (**Figure 2A1-L2, Suppl. Figure 3**) on juvenile maize leaves, covering different distances from the intercalary meristem (**Figure 2M**). This approach allowed us to reconstruct the morphogenesis of maize pavement cells using two consecutive time points. Our observations revealed that puzzle-like shapes in maize pavement cells emerge from a two-phase growth pattern: an initial phase of strong anisotropic expansion along the leaf’s longitudinal axis, followed by a shift to nearly isotropic growth, during which transverse expansion increases relative to longitudinal expansion. To simulate maize cell development, we initialized the model with a representative geometry of young cells (**Figure 2O**). By adjusting growth rates and anisotropy based on time-lapse observations, we reproduced the shapes and timing of maize puzzle cells (**Figure 2N**, **Figure 2P, Suppl. Movie 7**). We perturbed the model by reversing the growth sequence, initiating with transverse growth followed by longitudinal growth, while preserving the total amount of growth. This produced cell shapes that had lobes primarily oriented in the opposite direction, along the longitudinal axis of the leaf, different from those observed in maize (**Figure 2Q, Suppl. Movie 8-9**). These findings highlight the critical role of the dynamics of growth fields in determining cell shapes and explain the formation of cells with lobes that are primarily transversely oriented, even if the overall leaf shape is elongated. By integrating empirical observations with computational modeling, we show that the sequence of anisotropic and isotropic growth phases can explain the puzzle cell shapes observed in maize, demonstrating how temporal growth dynamics shape cell morphology *in planta*.

**Figure 2.**
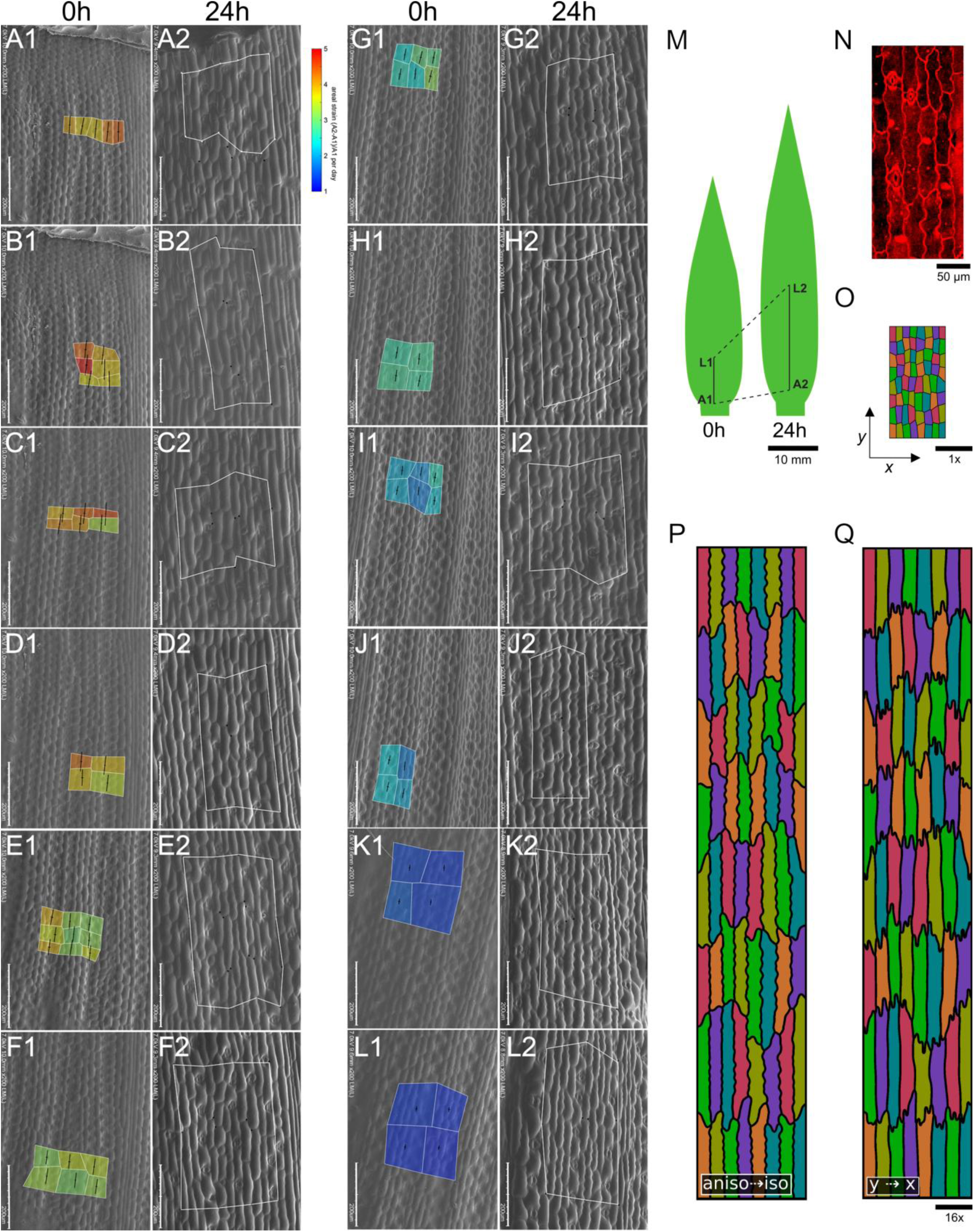
Temporal dynamics of maize (*Zea mays*) growth. (**A1-L2**) Electron micrographs show the temporal progression of maize growth and the relative cellular changes after 24 hours with images arranged sequentially along the leaf blade from the proximal base (**A**) to distal regions (**L**). In each paired set, such as **A1** and **A2**, area strain heat maps are presented, calculated as 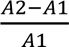, to visualize the daily growth rate for each region. Distinct boxes in the initial image of every sequence highlight this strain, and the lines within demonstrate the primary growth directions. The distances from the intercalary meristem on the specific blade regions are given: region **A** is in the meristematic region at 0h and at circa 200 μm at 24h; region **B** at 50 μm and 430 μm; region **C** at 185 μm and 1140 μm; region **D** at 320 μm and 1875 μm; region **E** at 620 μm and 3655 μm; region **F** at 760 μm and 4340 μm; region **G** at 960 μm and 5420 μm; region **H** at 1200 μm and 6630 μm; region **I** at 1500 μm and 7650 μm; region **J** at 1720 μm and 8650 μm; region **K** at 2400 μm and 10870 μm; region **L** at 3100 μm and 12730 μm. (**M**) Schematic drawings of maize leaf at 0h and 24h, indicating positions **A1** and **L1** on the 0h leaf and **A2** and **L2** on the 24h leaf. (**N**) Cell shape in the maize leaf epidermis under confocal microscopy. (**O**) Initial starting template of idealized maize cells. (**P)** Cell shapes that emerge when growth in the model initially follows anisotropic growth rates observed experimentally, then switches to more isotropic growth rates. (**Q**) The effect on ll shape when the template is first grown in the y direction, then the x direction to reach the same final template size as **I**. Scale bars, as indicated.

### Effects of disrupted organ growth on puzzle cell formation

Building on the maize results, we next examined how changes in overall organ growth patterns influence puzzle cell morphology. We asked whether growth in two directions, rather than strong elongation along one axis, predicts the development of lobes. As cells increase in area, they maintain low LEC values to minimize mechanical stress. Cells that expand in two directions reduce the LEC by developing lobes, whereas highly elongated, anisotropically growing cells can enlarge while maintaining small LECs and are therefore not expected to develop lobes. To quantify this, we introduced the concept of the *minimum axis* (min-axis), defined as the shortest dimension of the cell, measured by the width of the smallest rectangle that fully encloses it (**Figure 3A**). This measure better reflects the aspect of cell size relevant to mechanical stress; long, thin cells have low min-axis values, while cells that are large in both length and width exhibit higher min-axis values, which are expected to be positively correlated with lobeyness. Lobeyness is defined as the ratio of a cell perimeter to the perimeter of its convex hull, the inverse measure of convexity (**Figure 3A**). This measure is particularly effective at distinguishing lobey cells from long, thin cells with gentle curves but no pronounced lobes (*e.g.,* boomerang-shaped cells). By integrating the min-axis and lobeyness metrics, we provide a more robust framework for quantifying and distinguishing cell shapes, enabling better insights into the relationship between cell growth dynamics and mechanical stress. Our analysis of *Arabidopsis* revealed a stronger correlation between min-axis and lobeyness than between cell area and lobeyness (**Figures 3B-F**), consistent with the idea that combined min-axis and lobeyness better reflect mechanical constraints and stress-mitigation strategies.

**Figure 3.**
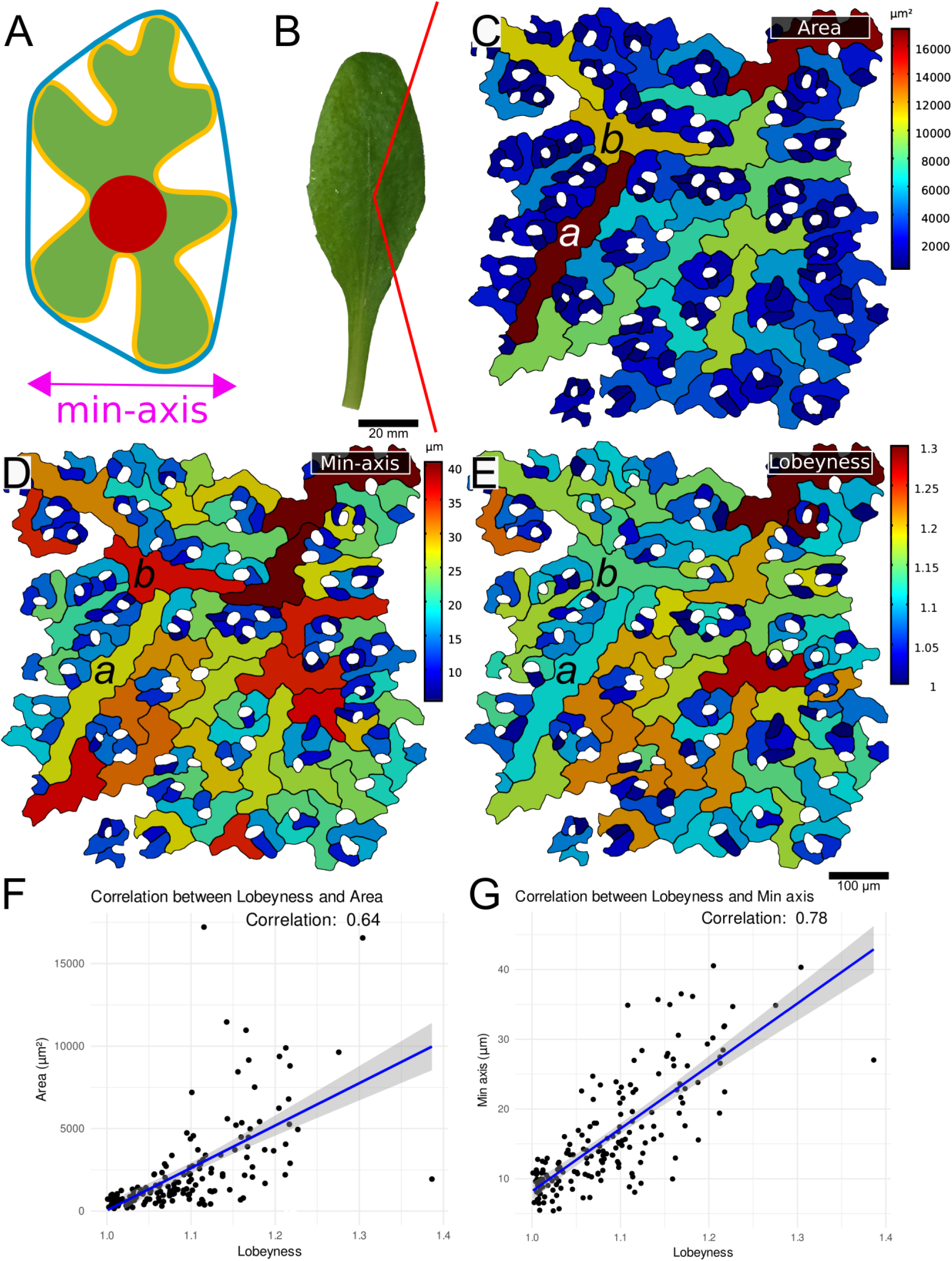
Lobeyness appears widely as a response to increasing cell size. (**A**) The visual depiction of the pavement cell descriptors: area is shown in green, lobeyness is defined as the cell perimeter (yellow) divided by the perimeter of the convex hull (blue), the min-axis (purple) is defined as the smallest width that would fit the cell, and the large open area (LEC) representing the magnitude of mechanical stress (red). (**B**) Image of a fully developed 3-week-old leaf of *Arabidopsis thaliana* Col-0. (**C-E**) Epidermal pavement cells from the leaf in (**B**), stained with propidium iodide, imaged with confocal microscopy, and segmented with MorphoGraphX. Cell templates are colored by area (**C**), min-axis (**D**), and lobeyness (**E**). (**F-G**) Graphs presenting the correlation between area and lobeyness (Corr = 0.64) (**F**) and between min-axis and lobeyness (**G**) for the same sample as in **C-E**. Scale bars, as indicated.

To explore how changes in organ shape and growth affect the shape of puzzle cells, we analyzed time-lapse imaging from 6 to 8 DAS in WT Col-0 and in the miR319-overexpressing *jaw-D* mutant (Harline et al., 2022), which downregulates TCP transcription factors, prolongs cell proliferation, and yields rippled leaves. We extracted multiple complementary metrics (**Figure 4**). Heatmaps of lateral (width) expansion ratios between 6–7 and 7–8 DAS (**Figure 4A–B**) show that *jaw-D* exhibits enhanced transverse growth compared to Col-0. Cell area heatmaps (**Figure 4C–D**) indicate that *jaw-D* cells remain smaller at all stages, and lobeyness heatmaps (**Figure 4E–F**) reveal consistently reduced puzzle-shaping in the mutant. Violin plots of log₂ lateral expansion ratios (**Figure 4G**) confirm increased width growth in *jaw-D*, while plots of log₂ cell area (**Figure 4H**) and lobeyness (**Figure 4I**) confirm that cells are smaller and less lobed. Across three biological replicates, linear mixed-effects analysis of lobeyness detected strong genotype, day, and genotype × day effects (all p < 0.001). Lobeyness was similar between genotypes at 6 DAS and 7 DAS, but significantly lower in *jaw-D* by 8 DAS. For cell area, *jaw-D* cells were significantly smaller than Col-0 at all stages (all p < 10⁻¹⁵). Proliferation heatmaps between 6–7 and 7–8 DAS (**Figure 4J–K**) show more frequent divisions in *jaw-D*, consistent with its higher total cell counts (**Figure 4M**). Mixed-effects analysis of cell number revealed a significant genotype × day interaction (p = 0.047), with counts diverging significantly only at 8 DAS. Accordingly, *jaw-D* leaves display a higher width-to-length ratio (**Figure 4L**), reflecting an altered balance between growth and division at the cellular level.

**Figure 4.**
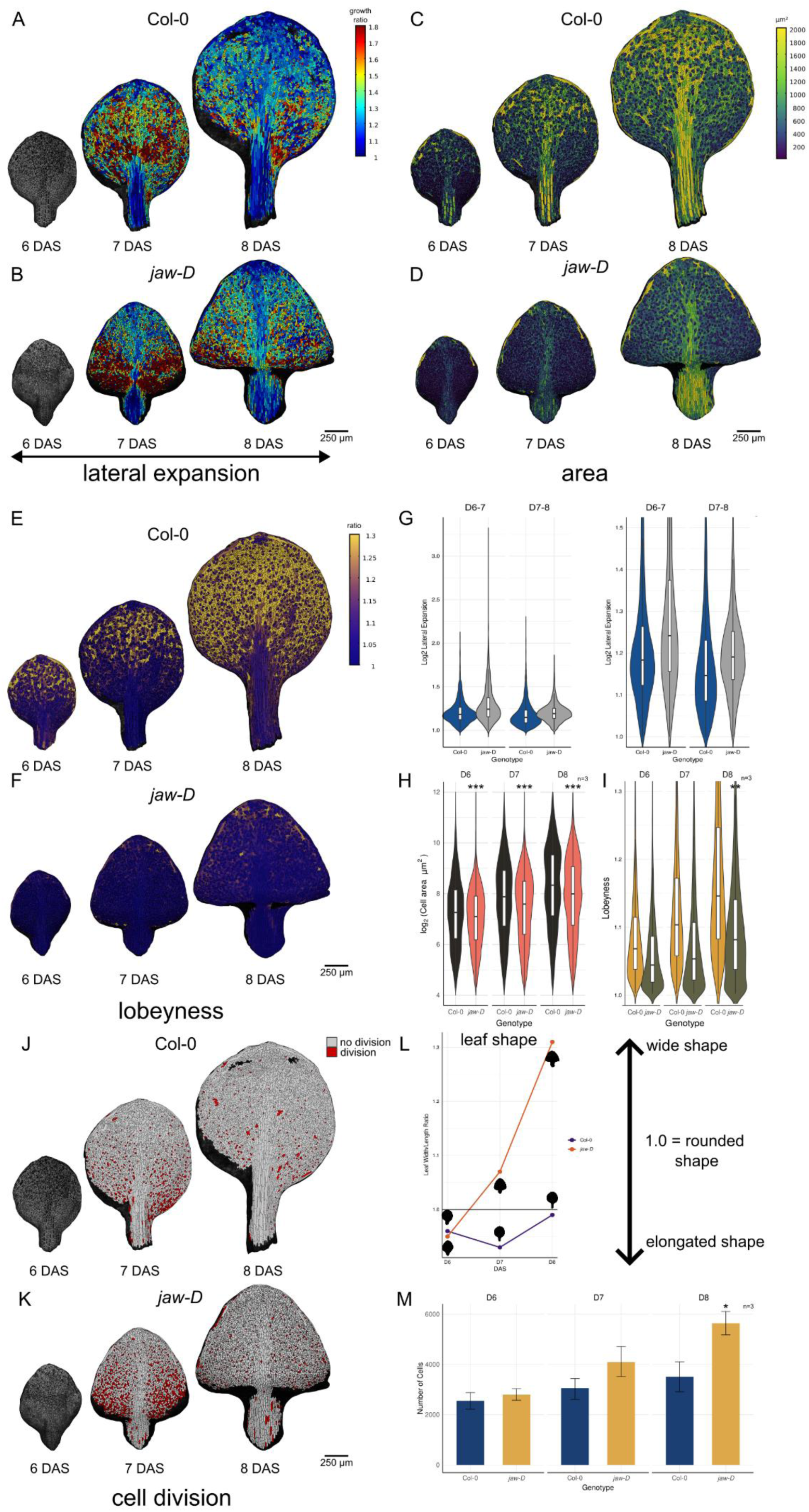
Cellular growth, shape and proliferation dynamics in Col-0 and *jaw-D* leaves. Data are derived from published time-lapse imaging of leaves in Col-0 and the miR319-overexpressing jaw-D mutant ^1^. (**A–B**) Heatmaps of lateral (width) expansion ratio between 6 and 7 days after sowing (DAS) and 7 and 8 DAS for Col-0 (**A**) and *jaw-D* (**B**), with warmer colors indicating greater relative width increase. (**C**–**D**) Cell area heatmaps at 6, 7, and 8 DAS for Col-0 (**C**) and *jaw-D* (**D**), color-coded by individual cell area. (**E**–**F**) Cell lobeyness heatmaps at 6, 7, and 8 DAS for Col-0 (**E**) and *jaw-D* (**F**), with warmer hues indicating higher lobeyness. **G**) Violin plots with internal boxplots of log₂ lateral expansion ratio measured for the samples shown in (**A**–**B**); full distributions are on the left and a zoomed inset on the right. (**H**) Violin plots with boxplots of log₂ cell area at 6, 7, and 8 DAS (n = 3 per genotype). (**I**) Violin plots with boxplots of cell lobeyness at 6, 7, and 8 DAS (n = 3 per genotype). (**J**–**K**) Proliferation heatmaps showing the number of cell divisions between 6 and 7 DAS and 7 and 8 DAS in Col-0 (**J**) and *jaw-D* (**K**); white indicates no division, red indicates one or more divisions. (**L**) Leaf width-to-length ratio for the samples in (**A**–**B**), where 1.0 corresponds to a circular outline, higher values represent wider shapes, and lower values indicate increasing elongation. (**M**) Bar plot of total cell number 3 per leaf (n = per genotype); error bars are ± SEM. Asterisks in (**H**), (**I**), and (**M**) indicate significance from linear mixed-effects model contrasts: *P* < 0.05 (\**), P < 0.01 (*), P < 0.001 (\*\*\**). *jaw-D* cells are smaller at all stages (all *P* < 10^-15^), show significantly reduced lobeyness only at 8 DAS (Col-0: 1.179 ± 0.001; *jaw-D*: 1.099 ± 0.001, *P* = 0.003), and divide more frequently, leading to higher total cell numbers at 8 DAS (Col-0: 1719 ± 40; *jaw-D*: 1912 ± 39, *P* = 0.016). Width-to-length ratios are consistently higher in *jaw-D*, reflecting altered growth balance. Scale bars, as indicated.

### Effects of disrupted puzzle cell formation on organ growth

We then asked whether the reverse relationship also holds by analyzing pavement cell morphology in mutants with defective puzzle cell formation and in wild-type plants subjected to pharmacological treatments. In mutants where puzzle shapes were absent, we did not observe cell rupture or death, as reported at the shoot apical meristem (Sapala et al., 2018). One possible explanation is that these cells experience lower mechanical stress, possibly due to changes in the arrangement of neighboring cells and tissue structure. Another hypothesis is that they have compensatory traits such as restricted organ growth, increased cell elongation, or altered cell wall properties. These may reflect developmental adjustments associated with the absence of puzzle-like lobes. Compared to the wild-type *Arabidopsis* (**Figure 5A, Suppl. Figure 4A**), the constitutively active Rho-of-Plants 2 (*CA-ROP2*) mutant, which alters actin filament accumulation, displayed non-lobed cells (Qiu et al., 2002). While lobeyness was abolished in *CA-ROP2*, the lack of lobes was partially compensated by elongated shapes (**Figure 5B, Suppl. Figure 4B**). Importantly, *CA-ROP2* mutants exhibited relatively mild reductions in overall plant growth, maintaining taller stature due to their more anisotropic growth habit, suggesting that puzzle cell formation inhibition in this line primarily impacts isotropically growing organs. Similarly, the *anisotropy1* (*any1*) mutant, which carries a mutation in cellulose synthase A1 (CesA1) affecting crystalline cellulose synthesis (Fujita et al., 2013), also lacked lobes but exhibited sinuous cells as a compensatory feature (**Figure 5C, Suppl. Figure 4C**). We analyzed the *constitutive triple response* (*ctr1*) mutant, a negative regulator of ethylene signaling, which displayed reduced cell expansion and lobeyness, with small cells and smaller leaves (**Figure 5D, Suppl. Figure 4D**). Collectively, these mutants indicate that reduced lobeyness is associated with altered cell shapes, such as elongation or zigzag morphologies. These changes could reflect responses to growth constraints but may also result from underlying disruptions in cell wall synthesis, anisotropy, or signaling pathways specific to each mutant background. Without direct functional evidence, the extent to which these altered shapes represent compensatory mechanisms remains unclear. Notably, all mutants with altered lobeyness exhibited lower correlations between min-axis and lobeyness, along with smaller leaves and overall reduced plant growth compared to wild type (**Figure 5E-H**). These observations reveal a consistent association between reduced lobeyness, small-er epidermal cells, and reduced plant growth. However, whether the loss of lobes contributes causally to these growth defects remains to be tested.

**Figure 5:**
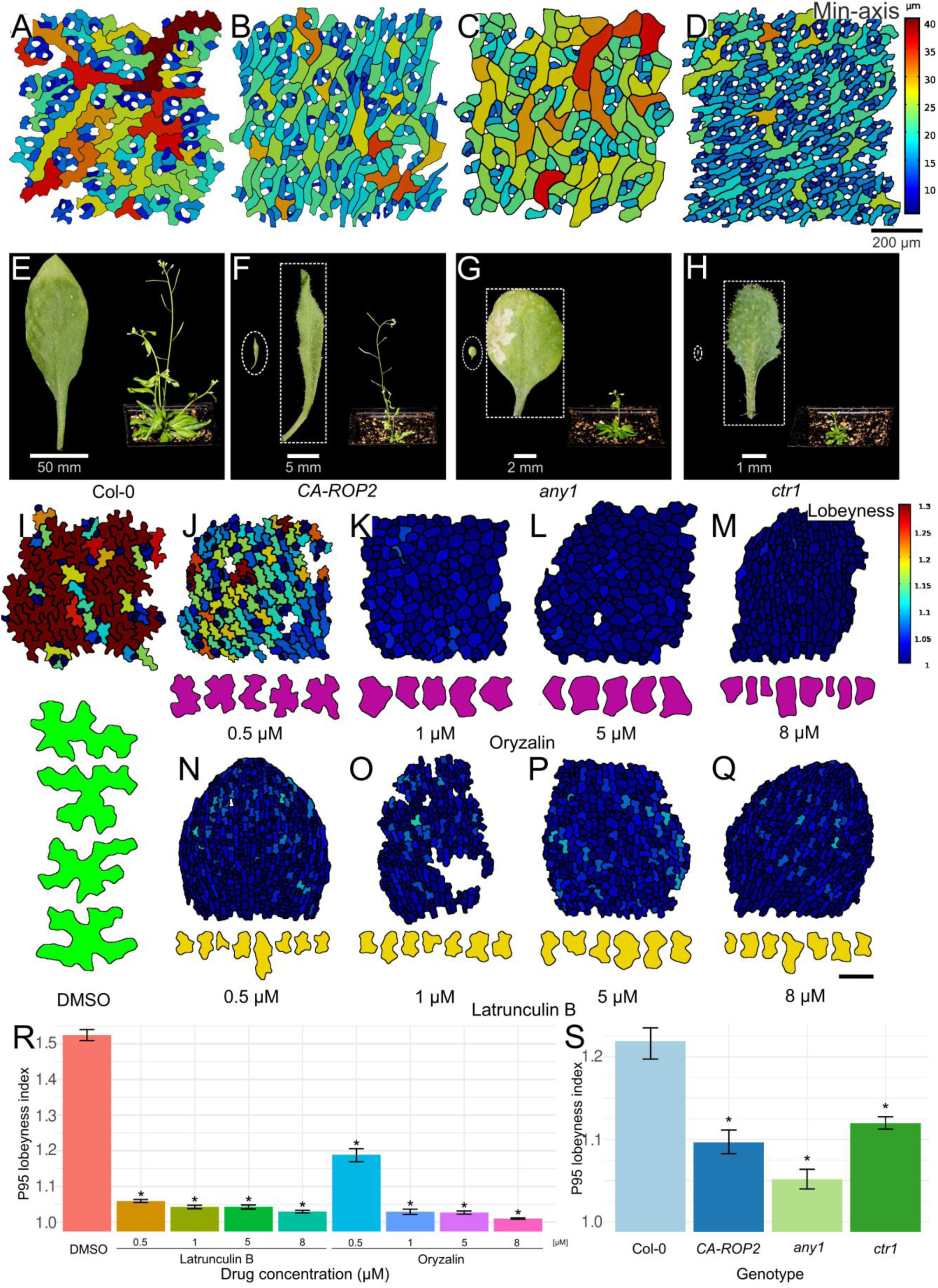
The lack of lobes is associated with growth defects. (**A-H**) The shape of epidermal pavement cells (**A-D**) and overall growth phenotypes (**E-H**) in 3-week-old *Arabidopsis thaliana* Col-0 and mutants. (**A**, **E**) Puzzle cells in Col-0 displaying high lobeyness and normal growth. (**B, F**) Thin and elongated epidermal pavement cells in the *CA-ROP2* mutant displaying decreased lobeyness (**B**), with long and thin leaves, and reduced overall plant growth (**F**). (**C, G**) Snaky epidermal pavement cells in the *any1* mutant displaying decreased lobeyness (**C**), with smaller and round leaves, and dwarf plant phenotypes (**G**). (**D, H**) Small epidermal pavement cells in the *ctr1* mutant displaying decreased lobeyness (**D**), with very small leaves, and dwarf plant phenotypes (**H**). (**E**-**H**) Each panel shows an image of the full leaf on the left and the pot with the full plant on the right; for mutants (**F**-**H**), the leaf within the dashed oval is scaled to match (**E**), while the dashed square provides a zoomed-in view with a separate scale bar. (**I-Q**) The effect of pharmacological treatments on the shape of epidermal pavement cells. (**I**) The mock treatment (DMSO) retains the typical morphology of puzzle cells. (**J-M**) Dose-dependent oryzalin treatments lead to a gradual decrease in lobeyness and an increase in LEC (cells labelled in violet). (**N-Q**) Dose-dependent latrunculin B treatments lead to a decrease in lobeyness with shallow lobes (cells labelled in yellow). (**R**) Treatments were administered at increasing concentrations of 0.5, 1, 5 and 8 µM. The 95th percentile of lobeyness is plotted for each drug treatment, with statistical significance determined using bootstrapping by pairwise comparisons to DMSO; significance is marked by asterisks and standard error. (**S**) The 95th percentile of lobeyness for different mutants, with statistical significance determined using bootstrapping by pairwise comparisons to Col-0; significance is marked by asterisks and standard errors. Scale bars: (**A**-**H)** as indicated on figure, (**I**-**Q**, contours) 50 µm, (**I-Q**, epidermis) 100 µm.

Having shown that reduced lobeyness in *Arabidopsis* mutants correlates with organ growth, we next used pharmacological treatments to acutely disrupt lobe formation in wild-type plants. This allowed us to test whether transient inhibition of lobing, independent of stable genetic mutations, produces similar effects. While maize was used to study how temporal changes in tissue-level anisotropy influence cell shape, *Arabidopsis* cotyledons expand nearly isotropically and are amenable to dose- and time-controlled perturbations of cytoskeletal components. Treatments with oryzalin, a microtubule depolymerizer, and latrunculin B, an actin polymerization inhibitor, were performed on cotyledons to examine the effects of inhibiting lobeyness. By employing dose- and time-dependent treatments, we precisely regulated the inhibitory effects. Compared to control samples (**Figure 5I**), low doses of oryzalin led to wider neck regions and larger LECs (**Figure 5J, K**). Higher doses further reduced lobeyness, and at very high concentrations, puzzle cell formation was completely abolished, resulting in nearly isodiametric cell shapes (**Figure 5L, M**). Latrunculin B treatments reduced lobeyness by reducing indentation depth, but even at high doses, lobe formation was not entirely eliminated (**Figure 5N-Q**), in contrast to oryzalin treatments. These treatments produced phenotypic responses similar to those observed in mutants, including reduced cotyledon sizes and overall inhibited plant growth. Although the perturbations used here broadly affect growth and cell wall dynamics in different ways, the consistency of phenotypic outcomes across independent manipulations supports the idea that reduced lobing contributes to the observed defects.

### Pavement cell shape reflects the developmental and environmental context of the plant

We have shown that epidermal pavement cell shape can depend on the growth dynamics of the organ in which they develop. This raises the question of how many plant species in previous analyses that did not have puzzle cells are actually able to make them in other developmental contexts with different growth. To address this question, we examined epidermal pavement cells across various developmental stages, organs, and environmental contexts. Pavement cells exhibit diverse shapes that vary not only across species but also among different organs within the same plant. Their morphology is influenced by developmental stage, environmental conditions, and tissue-specific cues, all of which alter growth dynamics and ultimately determine the extent of lobe formation (Liu et al., 2021; Majda et al., 2021; Zuch et al., 2022; Ikematsu et al., 2023). We first examined how developmental stage affects cell shape by comparing pavement cells in juvenile and mature leaves of various species. In quaking aspen (*Populus tremuloides*), juvenile leaves were less lobed than mature leaves (**Figure 6A, D**). A similar effect was observed in lilac (*Syringa vulgaris,* **Suppl. Figure 6A, B**) and camphor tree (*Camphora officinarum*, **Suppl. Figure 6C, D**). Notably, in lilac, cells previously reported as non-lobed (Vőfély et al., 2019) appeared lobed in our dataset, likely reflecting differences in developmental stage or environmental conditions. A likely explanation is that the timing of cell division cessation varied across species, with some plants delaying this process until later developmental stages. We frequently observed a coexistence of lobed and non-lobed cells within the same plant, or even within the same organ. For example, in elderberry (*Sambucus nigra*) leaves (**Suppl. Figure 7A**) and in the bracts of linden (*Tilia cordata*; **Suppl. Figure 7B**), cells situated above vascular tissues were elongated, while those above the mesophyll were lobed. Similarly, in early developing *Arabidopsis* leaves, the proliferative base contains isodiametric cells, with puzzle-shaped cells emerging distally in the more differentiated regions where division has stopped (Fox et al., 2018).

**Figure 6.**
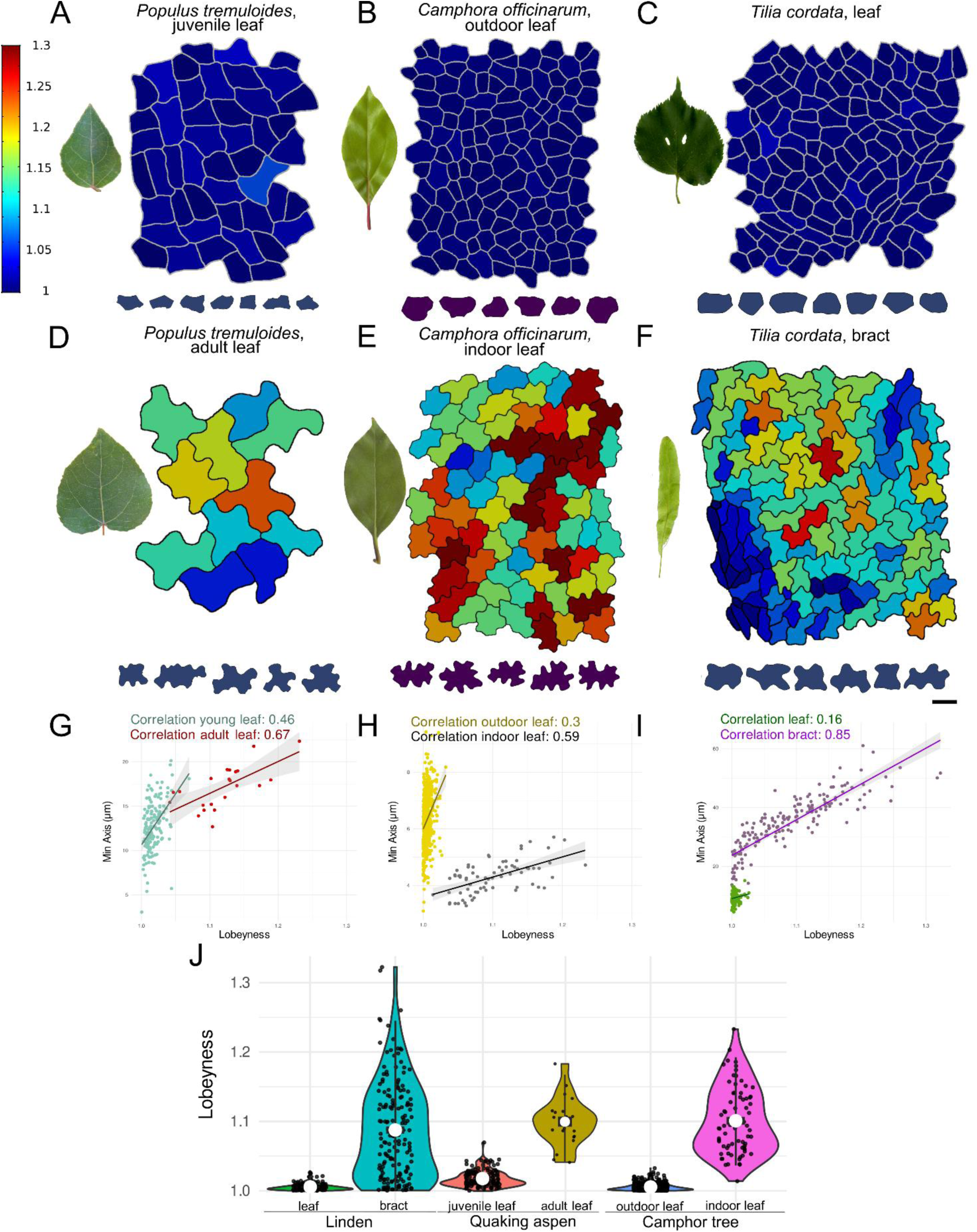
Pavement cell shapes exhibit developmental and environmental plasticity, varying across organs. (**A-F**) Leaf photographs are paired with lobeyness heatmaps of pavement cell contours and outlines near the 95th percentile of lobeyness. (**A**, **D**) Quaking aspen (*Populus tremuloides*), showing juvenile (**A**) and adult (**D**) leaves on the adaxial surface. (**B**, **E**) Camphor tree (*Camphora officinarum*) leaves grown outdoors (**B**) and indoors (**E**). (**C**, **F**) Linden (*Tilia cordata*) leaves (**C**) and bracts (**F**). (**G-I**) Scatter plots of the correlation between lobeyness and min-axis (the shortest cell dimension): (**G**) quaking aspen juvenile (Corr = 0.46) vs adult (Corr = 0.67) leaves, (**H**) camphor tree outdoor (Corr = 0.3) vs indoor (Corr = 0.59) leaves, and (**I**) linden leaf (Corr = 0.16) vs bract (Corr = 0.85). (**J**) Violin plots showing lobeyness distributions under each condition or organ, with the median indicated by a white circle, the interquartile range by a black box, and whiskers at 1.5 times the interquartile range. Scale bar: (**D** heatmap**, E** heatmap) 10 µm, (**A** heatmap) 15 µm, (**B, C, D** contours) 20 µm, (**B** heatmap, **E** contours) 25 µm, (**C** heatmap) 40 µm, (**F** heatmap, contours) 50 µm.

We found that pavement cell morphology often differs between the abaxial and adaxial surfaces of the same leaf. In species such as love-lies-bleeding (*Amaranthus caudatus*; **Suppl. Figure 8A-C**), Peruvian-lily (*Alstroemeria aurea*; **Suppl. Figure 8D-F**), fuchsia (*Fuchsia magellanica*; **Suppl. Figure 8G-I**), peppermint (*Mentha x piperita*; **Suppl. Figure 8J-L**), and cigar flower (*Cuphea ignea*; **Suppl. Figure 8M-O**), the abaxial surface shows more pronounced lobing than the adaxial surface, which is directly exposed to sunlight (Watson, 1942). In cigar flower, this may result from earlier cessation of cell division as the more lobed cells on the abaxial side are larger. In love-lies-bleeding, fuchsia, and peppermint, this size difference is absent, suggesting variation in the target LEC instead. These patterns highlight the influence of developmental context on cell morphology and indicate that model parameters such as target LEC size can substantially affect cell shape.

Environmental factors also significantly influence pavement cell shape. In camphor tree leaves, typically characterized by non-lobed cells in leaf epidermis (Zhao et al., 2006), cells developed puzzle shapes when grown in greenhouse conditions, with the min-axis-lobeyness correlation of 0.59, whereas this morphology was not observed when plants were cultivated outdoors with a correlation of 0.3 (**Figures 6B, E, H**). Outdoor-grown cells were larger, they remained non-lobed, while smaller indoor-grown cells adopted puzzle-like morphology, again suggesting a difference in target LEC size as a possible cause. The mechanism for how environmental signals could affect the LEC size is not clear, although the observations align with paleoecological studies that use cell sinuosity to distinguish sun and shade leaves and to infer environmental dynamics (Kürschner, 1997; Bush et al., 2017). The data also demonstrate that specific environmental conditions are another case where the appearance of puzzle-shaped cells may be suppressed, even though the species is able to make them.

Next, we wondered if plants that were reported not to make puzzle-shaped cells in leaves were able to make them in other organs. To this end, we examined pavement cell shapes in petals, sepals, and fruits. In certain species, such as linden (*Tilia cordata*; **Figure 6C**) and crownvetch (*Securigera varia*; **Suppl. Figure 9A**), the epidermal pavement cells on the adaxial side of mature leaves were non-lobed. However, despite the absence of lobes in leaves, lobed cells did appear in other organs, for example, in the bracts of linden (**Figure 6F**) or the petals of crownvetch (**Suppl. Figure 9B**). Interestingly, in linden, elongated bracts exhibited lobed cells, whereas rounder leaves did not (**Figure 6I**). Species without puzzle-shaped cells in leaves but with them in other organs were not rare (**Suppl. Figure 9C-L**), with puzzle cell shapes appearing in petals in cemetery iris (*Iris albicans*) and spring crocus (*Crocus vernus*), and sepals in St John’s wort (*Hypericum perforatum*), common gorse (*Ulex europaeus*) and blackthorn (*Prunus spinosa*). These results show that although these plants do not make puz-zle-shaped cells in leaves, the mechanism to make puzzle cells is nevertheless present.

### Prevalence of the mechanism to make puzzle-shaped cells

Because plant organs typically contain a mixture of lobed and non-lobed cells, the average cell shape may not capture the capacity of a species to produce puzzle-shaped cells. To address this, we determined the ability of each species to develop lobed cells by using the 95th percentile of cell lobeyness, which represents the upper range of lobing across different organs and developmental contexts (**Figure 6B, C**). The rationale is that if the plant can make lobed cells, it must possess the mechanism, even if most of its cells are non-lobed. This metric revealed that most species possess the capacity to generate highly lobed cells, even if the average cell shape does not exhibit significant lobeyness as previously reported by Vőfély et al. (2019).

To assess the universality of lobe formation in response to increasing cell size, we conducted a large-scale analysis of the correlation between the min-axis and lobeyness across vascular plant species (**Figure 7A**). We analyzed cell shapes in 327 species and 663 species-organ combinations, totaling 72,026 cells (**Suppl. Table 1**). In our analysis, we collected samples from 45 species in various plant organs and incorporated data from previously published analysis of leaf epidermal pavement cells in 19 species Sapala et al. (2018), 250 species from Vőfély et al. (2019), and data from 13 early Miocene species (Reichgelt et al., 2020). In 235 of 327 species (72%), we observed a significant positive correlation (≥ 0.3) between min-axis and lobeyness, with particularly high correlations found in tobacco (*Nicotiana tabacum*, Corr = 0.96), cigar plant (*Cuphea ignea*, Corr = 0.90), morning glory (*Ipomoea tricolor*, Corr = 0.91), black nightshade (*Solanum nigrum*, Corr = 0.90), Java fern (*Microsorum pteropus*, Corr = 0.85), and Peruvian lily (*Alstroemeria aurea*, Corr = 0.89) (**Suppl. Figure 10A–F**, **Suppl. Figure 11A**). Overall, these findings support the hypothesis that the puzzle cell shape is a widespread adaptive strategy for managing mechanical stress during growth.

**Figure 7.**
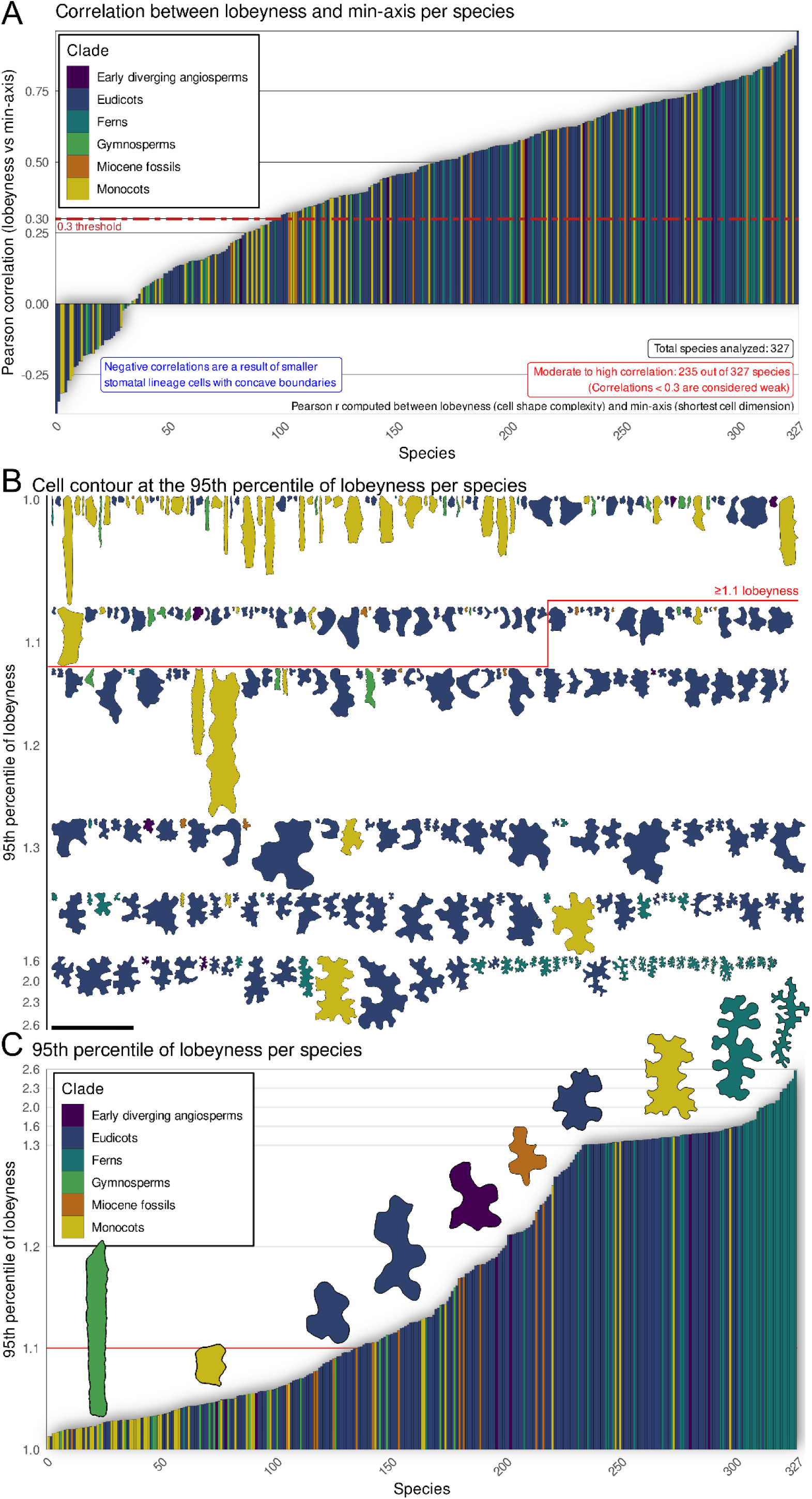
Lobeyness is a common feature observed across different species. **(A)** Bar plot displaying the Pearson correlation (r) computed between lobeyness (cell shape complexity) and min-axis (the shortest cell dimension) across 327 species. Correlations below 0.3 are deemed weak while those at or above 0.3 are classified as moderate to high. In our analysis, 235 out of 327 species exhibited moderate to high correlations. The 0.3 threshold was selected as a heuristic cutoff to differentiate species with pronounced positive relationships from those with less pronounced associations. Bars are color-coded by clade. (**B**) Visual representation of a single pavement cell contour per species, selected at the 95th percentile of lobeyness. The contours are arranged in order of increasing cell shape complexity, color-coded by clade. The 1.1 threshold was selected as a heuristic cutoff to differentiate species with significant lobeyness from those with less pronounced lobes. (**C**) Bar plot of the 95th percentile of lobeyness values with selected cell contours superimposed. The contours are color-coded by clade. The contours at approximate lobeyness values 1 (mountain cypress, *Widdringtonia nodiflora*), 1.05 (Panama hat plant, *Carludovica palmata*), 1.1 (yellow trumpetbush, *Tecoma stans*), 1.15 (garlic mustard, *Alliaria petiolata*), 1.2 (Japanese star anise, *Illicium anisatum*), 1.25 (colicwood, *Myrsine*), 1.3 (yarrow, *Achillea millefolium*), 1.5 (Peruvian lily, *Alstroemeria aurea*), 2 (*Phanerophlebia*), and 2.5 (rough maidenhair fern, *Adiantum hispidulum*) were selected to represent all clades at increasing positions along the lobeyness gradient (minimum 1.0 to maximum 2.6). Emphasis is given to the range from 1.0 to 1.3, as significant lobeyness begins to appear around a lobeyness value of 1.1. Scale bars: (**C** contours lobeyness values 1, 1.05, 1.15, 1.25, 1.3, 2, 2.5) 100 µm, (**C** contours lobeyness values 1.1, 1.2) 200 µm, (**C** contours lobeyness value 1.5) 400 µm, (**B**) 500 µm.

However, some species displayed weak or even negative correlations between the min-axis and lobeyness (**Figure 7A**), which seems contradictory to the notion that lobes form in response to a developmental constraint based on stress. Upon further examination of these cases, we identified several reasons for the lack of a strong correlation in certain samples. In cardboard palm (*Zamia furfuracea*), the highly anisotropic growth led to elongated cells without an increase in the min-axis, thus not favoring lobe formation (**Suppl. Figure 10H, Suppl. Figure 11B**). Conversely, species like yellow guava (*Psidium guajava*) and pinwheel flower (*Tabernaemontana divaricata*) maintained a small uniform cell size, suggesting frequent divisions to maintain size, even though the leaves can become quite large (**Suppl. Figure 10G, J, Suppl. Figure 11C, D**). Weak correlations were also seen in some species with highly lobey cells, such as hare’s foot fern (*Davallia solida*; **Suppl. Figure 10I, Suppl. Figure 11E**). The min-axis does not capture this case well, as substantial variability comes from branching rather than an increase in lobes, as the cells are all highly lobey. Here, it would be better to sample a patch of the leaf where the cells have a gradient from smaller dividing cells to mature expanded ones. In species without lobes, like agapanthus (*Agapanthus praecox*), cell staggering had a greater impact on thinner, less elongated cells (**Suppl. Figure 10L, Suppl. Figure 11F**), whereas in Virginia spiderwort (*Tradescantia virginiana*), irregular shapes in smaller stomatal lineage cells increased the tissue’s lobeyness scores (**Suppl. Figure 10K, Suppl. Figure 11G**). Overall, our analysis shows that species that made lobed cells had significant positive correlation between lobeyness and min-axis, and that species where this correlation was absent largely fall into a small number of well-defined classes.

We hypothesize that puzzle cell shape reduces stress enlarging cells by controlling LEC size. Consequently, large cells, compared to small cells, are expected to exhibit a smaller increase in LEC for the same increase in area. To test this prediction, we plotted cell area vs LEC and used a quadratic fit line passing through the origin (**Suppl. Figure 12A**). If our hypothesis holds, the curve should bend to the right (*α*< 0), indicating that the LEC area increases more slowly than cell area. We observed exactly this pattern (**Suppl. Figure 12B, C**). Moreover, since uncontrolled expansion in all directions would proportionally increase LEC, cells must preferentially become either elongated or lobed. Using the min-axis as a proxy for directional growth, we found that LEC area increases more slowly than the min-axis (**Suppl. Figure 12D, E**), a relationship that persisted across multiple clades and species (**Suppl. Figure 12F-K**). Overall, 87% of species (286 of 386) exhibited negative α values (**Suppl. Figure 13**). An exact sign-test confirms that this pattern is highly unlikely to occur by chance (p < 0.05), indicating that LEC area increases more slowly than the min-axis. This finding reveals a consistent association between increased lobing and reduced LEC expansion relative to growth, supporting the idea that lobing may help to limit wall free span and reduce tensile stress. However, whether these shapes actively buffer stress or simply arise in response to mechanical constraints remains to be tested.

## Discussion

Since the epidermis plays a pivotal role in shaping plant organ architecture (Kutschera and Niklas, 2007; Beauzamy et al., 2015), it is essential that its cells possess robust mechanisms to mitigate mechanical stress. Here we show that puzzle cell shapes emerge as a result of the tight interaction between mechanical constraints and growth at the organ scale. Our analysis of epidermal pavement cells in maize leaves and computational modeling underscore the critical role of specific growth trajectories in shaping puzzle cells, suggesting that the morphology of these cells may offer insights into the growth history and environmental adaptations of both living and extinct plant species (Couturier et al., 2009). In many cases, the observed variation in the presence or absence of puzzle cells across different contexts likely reflects differences in growth and proliferation dynamics rather than fundamental differences in the mechanism of puzzle cell patterning. In *jaw-D*, time-lapse heatmaps reveal a clear increase in transverse (lateral) expansion, concurrent with smaller, less-lobed cells and elevated division rates; these cellular dynamics map precisely onto the broader, more triangular leaf form. Thus, the puzzle shape phenotype in *jaw-D*, it is more likely related to its effect on cell proliferation and growth, rather than involvement in puzzle cell patterning directly. This implies that when studying genes that have small to moderate effects on puzzle cell shape, it is necessary to consider that any gene influencing growth rate, timing, directionality, or cell division would be expected to influence puzzle cell shape. Differences in puzzle cell shape could reflect growth differences, or differences in cell division timing, and must be excluded before proposing a role in the lobe patterning process itself (Xu et al., 2010).

Genetic modifications in *Arabidopsis* that affect growth anisotropy have puzzle cell phenotypes. Examples are the *ftsh4* mutant that makes sepals nearly isotropic due to altered ROS levels. In this mutant the increased isotropy of growth causes giant sepal cells to adopt puzzle shapes. Conversely, *LONGIFOLIA1* overexpression leads to long, thin leaves with elongated cells and reduced lobing (Wang et al., 2016; Sapala et al., 2018). Genes that promote cell proliferation lead to the formation of smaller, less lobed cells, as observed in the *lgo-2* mutant (**Suppl. Figure 5B, D**). On the contrary, genes that inhibit cell division and promote cell expansion, such as those involved in endoreduplication, result in larger, more lobed cells, as in the *LGO OX* line (**Suppl. Figure 5C, D**). Interestingly, recent work reports a correlation between local wall stress levels and lobe width in puzzle cells, raising the possibility that mechanical forces contribute to shaping lobes (Malivert et al., 2021). This intertwining of growth dynamics and cell proliferation with the development of puzzle cell shapes highlights the importance of distinguishing between genes specifically affecting the mechanisms of puzzle cell formation from those controlling growth amount, growth anisotropy, and cell division.

Although we cannot definitively claim broad conservation based solely on these experiments, our findings are consistent with the idea that the underlying mechanism involves microtu-bule-guided cellulose deposition, which restricts growth in localized regions of the cell wall. Pharmacological treatments using oryzalin and latrunculin B further support this model. Oryzalin, which depolymerizes microtubules, disrupts localized cell wall reinforcement in the neck regions of developing puzzle cells, while latrunculin B, by inhibiting actin polymerization, impairs the targeted delivery of key cell wall components (Qiu et al., 2002; Panteris and Galatis, 2005; Bidhendi et al., 2019). Together, these results confirm the Sapala et al. (2018) model prediction that puzzle cell formation relies on regulated growth restriction mediated by cellulose microfibrils and associated mechanical ‘springs’, rather than by unrestrained cellular outgrowths. *Arabidopsis* mutants unable to form puzzle cells exhibit stunted growth, underscoring the importance of the shape to enable cells to expand for normal development. While some plants developed snaky cells as a compensatory response to the absence of puzzle cells, this alternative morphology does not fully restore normal cell formation and is rarely ob-served in both extant and fossil records, suggesting that it may be evolutionarily disfavored compared to the more common and adaptable puzzle cell morphology. Collectively, these observations support the view that puzzle cell formation emerges as a developmental response to mitigate the high mechanical stresses that would be created in large isodiametric cells, and that it is mediated by growth restriction, a process that is likely broadly conserved in vascular plants.

Despite the apparent complexity of factors influencing puzzle cell formation, their wide-spread emergence reflects a response to a physical constraint shared by all plants. Puzzle cell formation is essential in organs with large, isotropically growing cells to prevent the formation of extensive, unreinforced areas that would bulge out under high mechanical stress. However, puzzle cell shape is only required when cells become large in more than one direction, and multiple strategies may mitigate this need. For example, some species, such as yellow guava (*Psidium guajava*), appear to maintain smaller, more isodiametric epidermal cells, potentially through higher division rates, which would limit cell enlargement and thus lessen the mechanical demand for lobing. Alternatively, many plants transition from actively dividing, meristematic cells to larger, endoreduplicated cells (**Suppl. Figure 5**), a process common in the epidermis of leaves, roots, and sepals of *Arabidopsis,* that often promotes puzzle cell formation as cells expand. In roots and hypocotyls, growth is highly anisotropic resulting in large but elongated cells; in these cases, the elongated shape minimizes open areas and makes puzzle cell formation unnecessary. However, in organs where growth is less anisotropic, such as leaves and cotyledons, larger cells adopt the puzzle shape to mitigate bulging and mechanical stress. Under isotropic growth conditions, the absence of puzzle cells would lead to the development of large, unreinforced regions in the periclinal cell wall, exposing cells to considerable turgor-induced stress. Another strategy to manage stress in large, isodiametric cells would be to develop thicker cell walls; however, this approach is uncommon, likely because it requires substantially more resources than forming a cell shape with lobes and indentations (Majda et al., 2017; Majda et al., 2019). Interactions with subepidermal tissues may also contribute to epidermal lobing. The mesophyll contains air spaces that alter local mechanical support, and differences in tissue curvature between adaxial and abaxial surfaces could influence the distribution of wall stress. Such factors may help explain why lobing differs between the two surfaces in *Arabidopsis* and other species. Although direct experimental evidence remains limited, these interactions represent an important frontier for understanding how epidermal and subepidermal layers coordinate during organ growth. Our large-scale analysis of pavement cells across vascular plants, including plant organs beyond leaves such as sepals, bracts, and even fossil species (Reichgelt et al., 2020), showed that most species contain cells that exhibit significant puzzle cell shapes (**Figure 7**), whereas species lacking puzzle cells tend to have elongated cells or cells with small sizes. Interestingly, we found that while some species may not exhibit puzzle cells in their leaves, they can still form them in other organs, such as sepals or bracts, and that the appearance of puzzle cells is influenced by developmental stage and environmental conditions. Overall, our integrated analysis reveals that puzzle cell formation is a dynamic, evolutionarily conserved strategy that enables plants to optimize cell and tissue integrity in response to mechanical constraints, thereby ensuring robust organ development across diverse environmental contexts.

## Author contribution

**Conceptualization:** RSS, MM. **Methodology:** NT, RSS, MM. **Investigation:** NT, BL, AP, FC, LH, AM, TR, DK, AR, RSS, MM. **Formal analysis:** NT, FC, DK, AHKR, AR, RSS, MM. **Data curation:** NT, RSS, MM. **Writing – original draft:** MM. **Writing – review & editing:** NT, BL, TR, DK, AHKR, AR, RSS, MM**. Funding acquisition:** NT, AHKR, DK, AR, RSS, MM. **Supervision**: AHKR, RSS, MM.

## Acknowledgements

We are grateful to Olivier Hamant, Arezki Boudaoud and Alice Malivert for valuable comments on the manuscript. We thank John Innes Foundation (NT), the University of Lausanne (NT, AP, LH, AM, MM), EMBO Postdoctoral Fellowship (LH), NSF IOS 1553030 (AHKR), the National Science Centre, Poland (SHENG1 grant 2018/30/Q/NZ3/00189) (DK), ERA-CAPS (DK, RSS, MM), the Sciences and Engineering Research Council of Canada Discovery Grant 2021-02795 (AR), and the Biotechnological and Biological Sciences Research Council (BBSRC) Institute Strategic Programme Grant (BB/X01102X/1) to the John Innes Centre (RSS). We acknowledge Dr Izabela Potocka from the Scanning Electron Microscopy Laboratory of Institute of Biology, Biotechnology and Environmental Protection, University of Silesia for help in SEM of replicas. We thank Dr Azahara Martin for providing us with access to the phase-contrast microscope. We would like to thank Prof Geoffrey Wasteneys for providing *any1* seeds, and the Nottingham Arabidopsis Stock Center for distributing other seeds. The confocal microscopy was performed at the Cellular Imaging Facility (CIF), Uni-versity of Lausanne.

