## Supplemental Information for "Growth history leaves a geometric trace in puzzle cells"

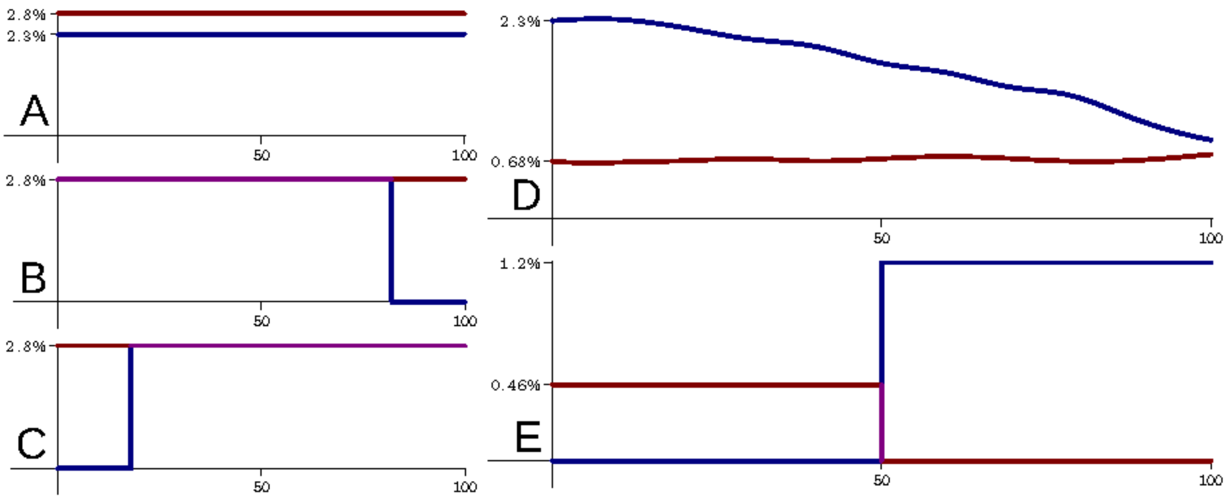


**Suppl. Figure 1. Growth rates used in simulations shown in Figure 1-2.** All simulations ran for 100-time steps. Graphs show relative expansion per time step in X (blue) and Y (red) directions. (**A–C**) Total growth was 10-fold in the X direction and 16-fold in the Y direction. (**A**) Uniform anisotropic growth (Figure 1D, G). Growth rates are constant over time, with growth in Y slightly higher than in X. (**B**) Isotropic growth followed by anisotropic growth (Figure 1E, H). Growth in Y is constant over the entire growth period. Growth in X matches it until the final width is reached, at which point growth in X ceases. (**C**) Anisotropic growth followed by isotropic growth (Figure 1F, I). The same as **B**, but the X growth period comes at the end of the simulation rather than the beginning. (**D–E**) Total growth was 6-fold in the X direction and 2-fold in the Y direction. (**D**) Growth rates drawn from measurements of maize (Figure 2P). The growth rate in the Y dimension remains stable. The growth rate in X starts high and then decreases overtime. (**E**) Maize template with very different growth rates (Figure 2Q). All growth in Y occurs in the first half of the simulation, while all growth in X occurs in the second half.

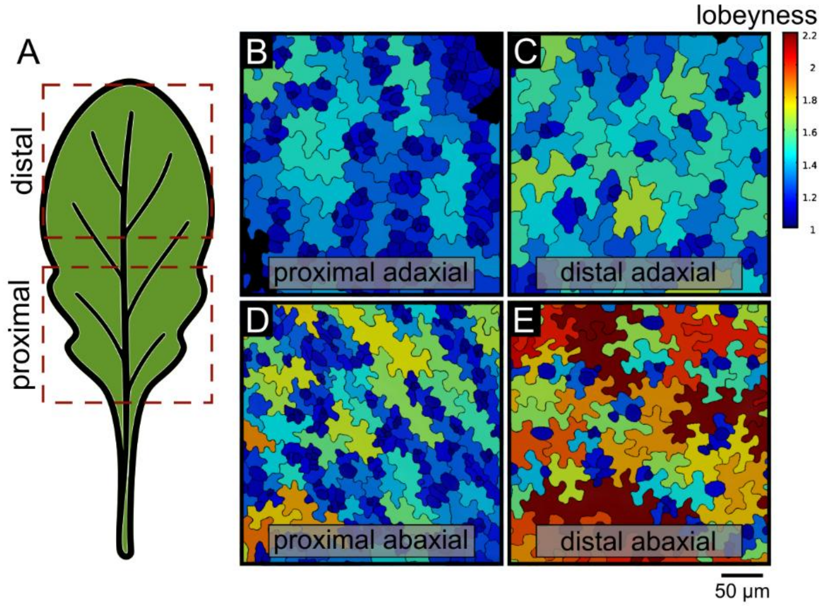


**Suppl. Figure 2. Regional variation in pavement cell lobeyness across *Arabidopsis* leaf. (A**) Schematic of a mature Arabidopsis leaf showing red dashed boxes demarcating the proximal and distal sampling regions on both the adaxial (upper) and abaxial (lower) surfaces. (**B–E**) Lobeyness heat-maps of segmented pavement cells in each indicated region: (**B**) proximal adaxial, (**C**) distal adaxial, (**D**) proximal abaxial, (**E**) distal abaxial. In each panel, individual cells are color-coded by lobeyness (warmer colors = higher lobeyness), illustrating that lobing is more pronounced in distal versus proximal regions and more elevated on the abaxial surface compared to the adaxial surface. Scale bars, as indicated.

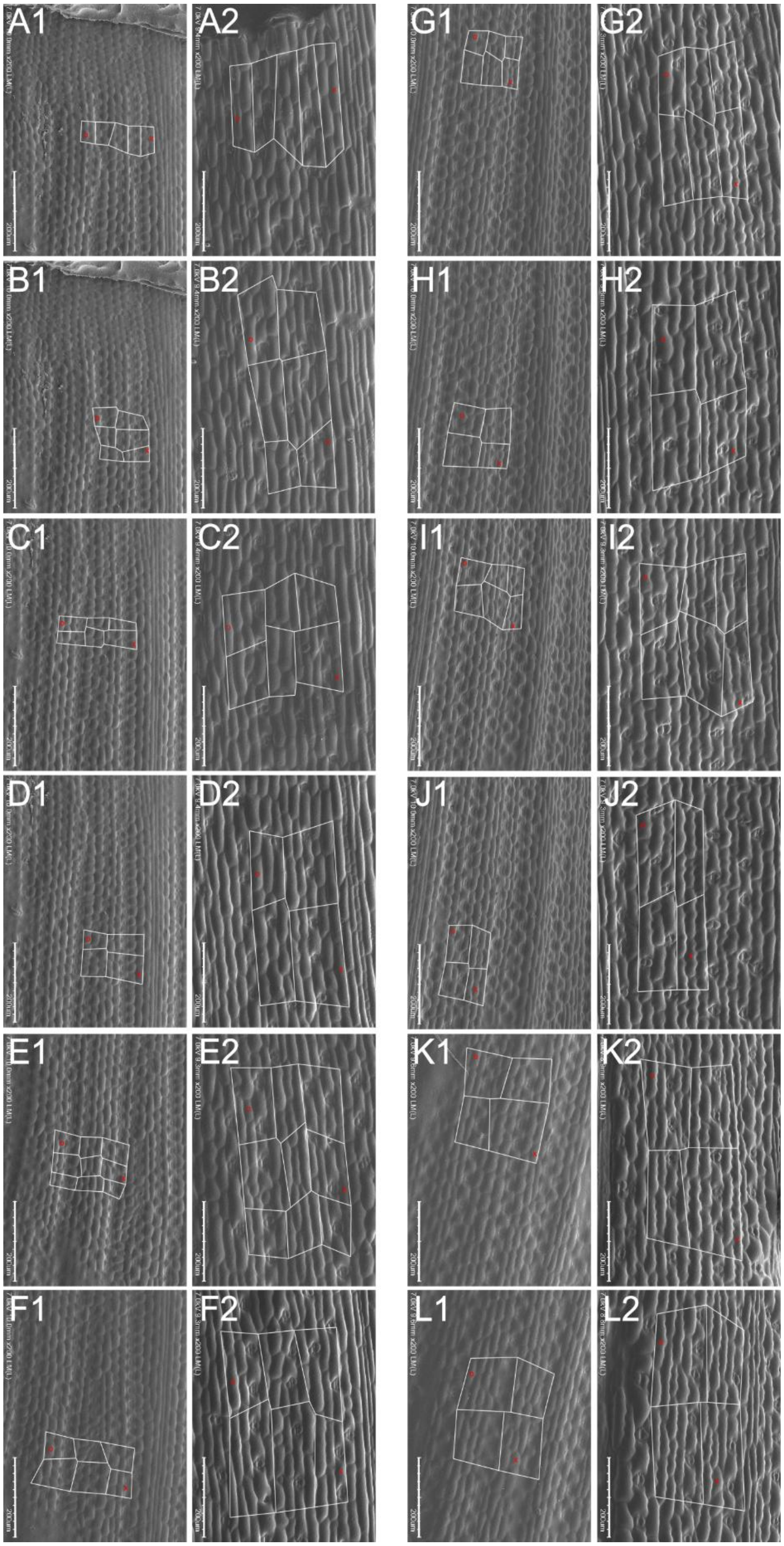


**Suppl. Figure 3. Landmarks and cellular tracking in maize growth. (A1-L2)** Electron micrographs, as in Figure 2, with landmarks used to calculate growth shown. Specific cells are marked with red O or X in each paired set (*e.g.*, A1 and A2) to track individual cells at 0h and 24h. Scale bars, as indicated.

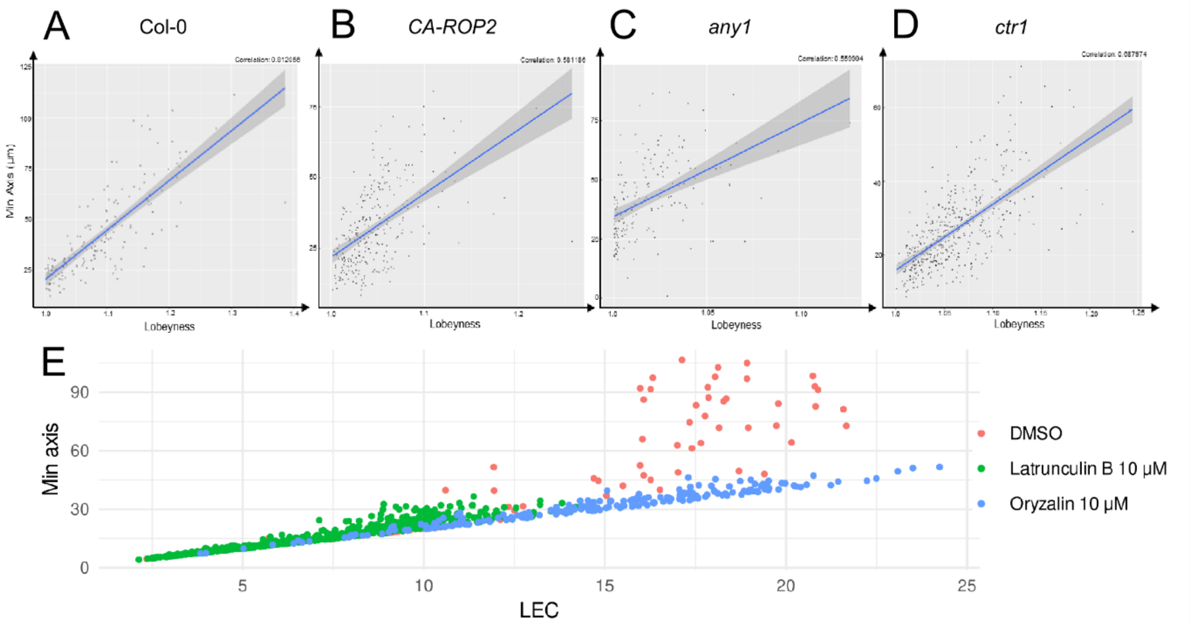


**Suppl. Figure 4. Correlations between the min axis and lobeyness in various mutants and with LEC in drug-treated plants.** (**A**) Puzzle cells in the wild type exhibit a pronounced correlation between min axis and lobeyness (Corr = 0.81). (**B-D**) This correlation is notably diminished in mutants: *CA-ROP2* (Corr = 0.58) (**B**), *any1* (Corr = 0.56) (**C**), and *ctr1* (Corr = 0.69) (**D**). (**E**) The scatter plot reveals the relationship between min axis and LEC in pavement cells of drug-treated plants: DMSO control, latrunculin B 10 μM, and oryzalin 10 μM. The nearly linear trend in drug-treated plants suggests a diminished or absent capacity for these plants to form lobes.

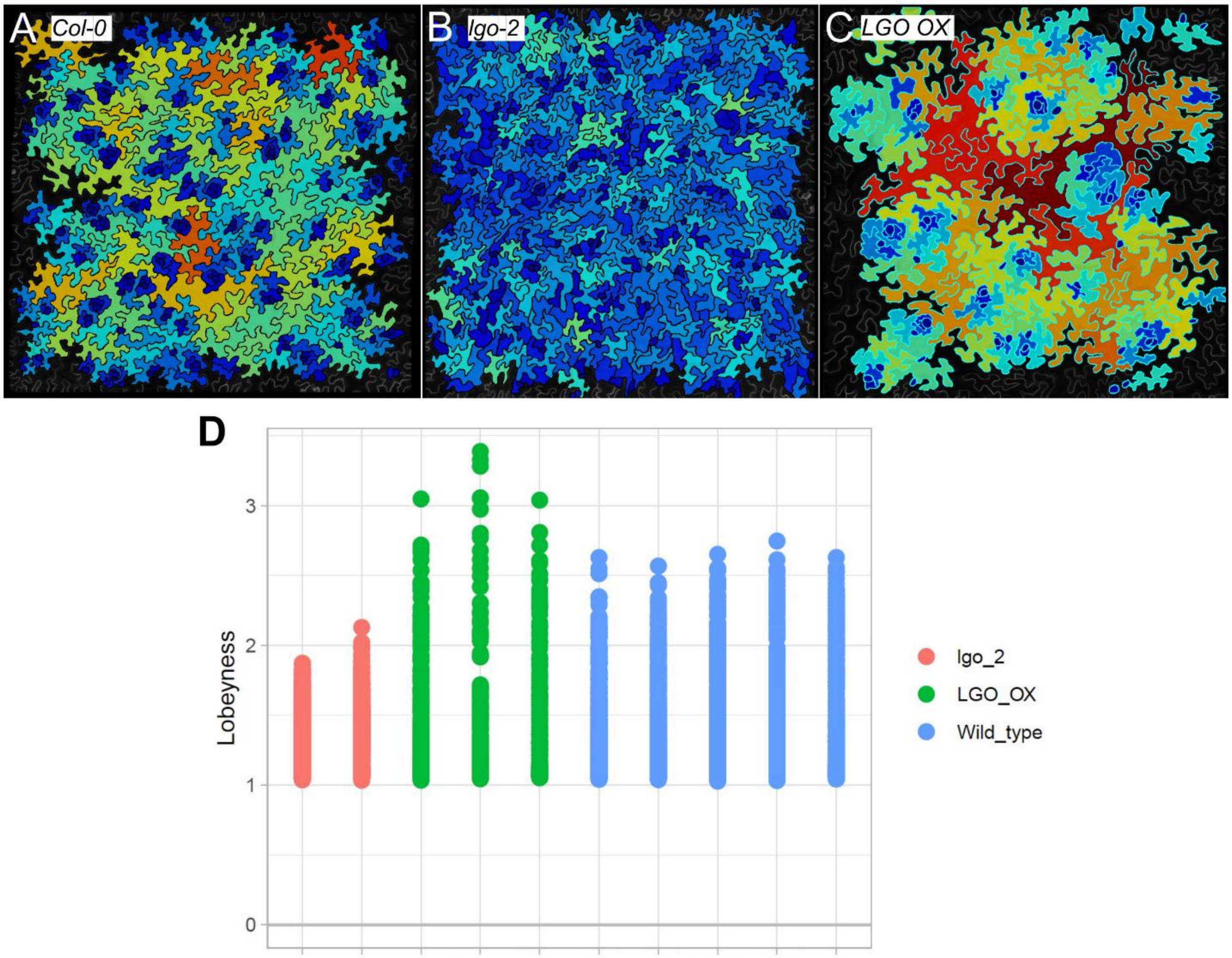


**Suppl. Figure 5. Lobeyness of pavement cells is negatively correlated with cell division activity.** (**A-C**) Heat maps represent the lobeyness of pavement cells in the wild type (**A**), *lgo-2* mutant (**B**) and *LGO-OX* (**C**) on abaxial side of the leaf in *Arabidopsis* 25 days post germination. (**D**) Quantification of lobeyness in the wild type, *lgo-2* mutant, and *LGO-OX*. Reduced lobeyness is observed in the *lgo-2* mutant compared to wild type, while increased lobeyness is seen in the *LGO-OX*, indicating a negative correlation between pavement cell lobeyness and cell division activity regulated by *LGO*.

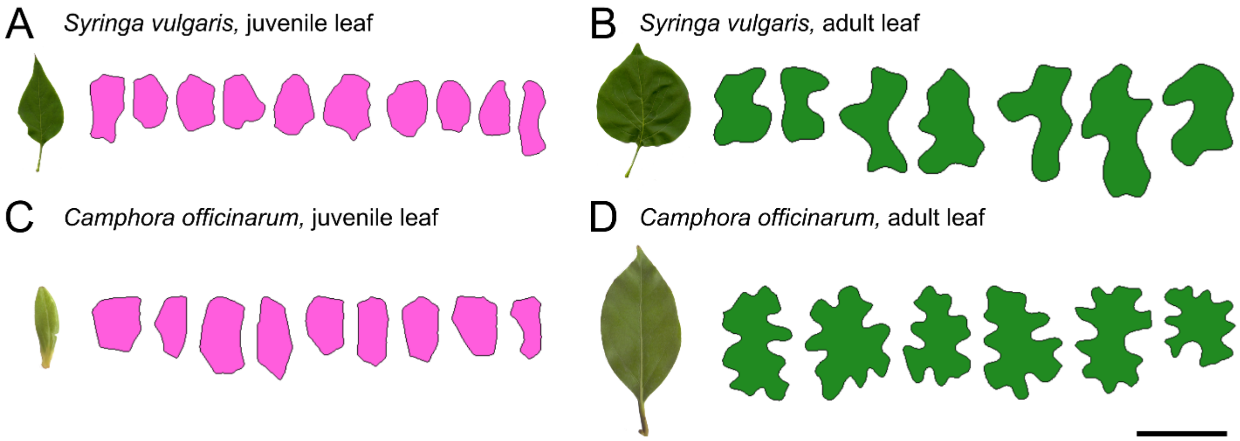


**Suppl. Figure 6. Developmental variation in leaf pavement cell shapes.** (**A-D**) Juvenile leaf and corresponding cell outlines (**A**) compared to adult leaf with more lobed cells (**B**) in lilac (*Syringa vulgaris*). Juvenile leaf and corresponding non-lobed cell outlines (**C**) compared to adult leaf with lobed cells (**D**) in camphor tree (*Camphora officinarum*) grown indoors. Different colors indicate cells sampled from: small leaves (pink), adult leaves (dark green). Scale bars for cell contours: 50 µm (**A**), 75 µm (**C, D**), 200 µm (**B**).

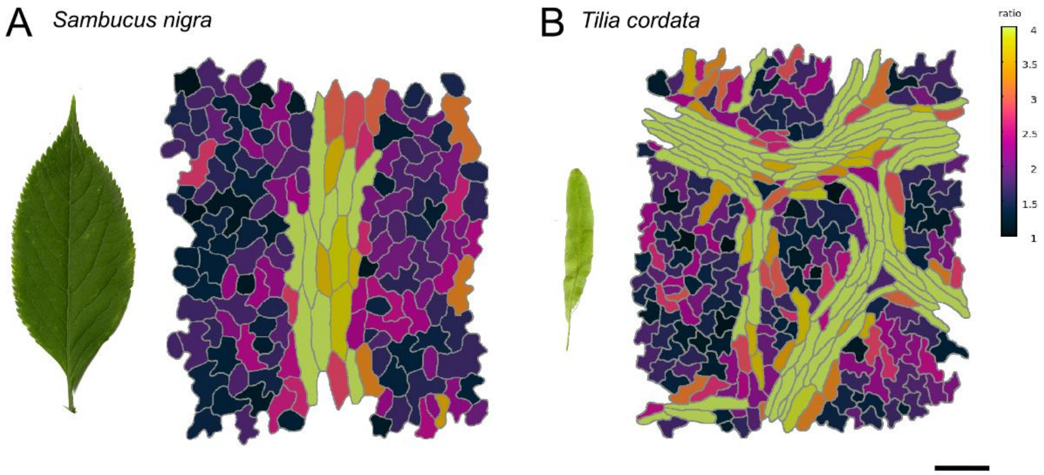


**Suppl. Figure 7. Variation in epidermal pavement cell morphology on the organ surface.** (**A**) Elderberry (*Sambucus nigra*) leaf and corresponding contours of the pavement cell on its abaxial side. (**B**) Linden (*Tilia cordata*) bract and the corresponding contours of the pavement cell on its surface facing the flower. Cells are color-coded by aspect ratio heatmap. Scale bars for cell contours: 100 µm.

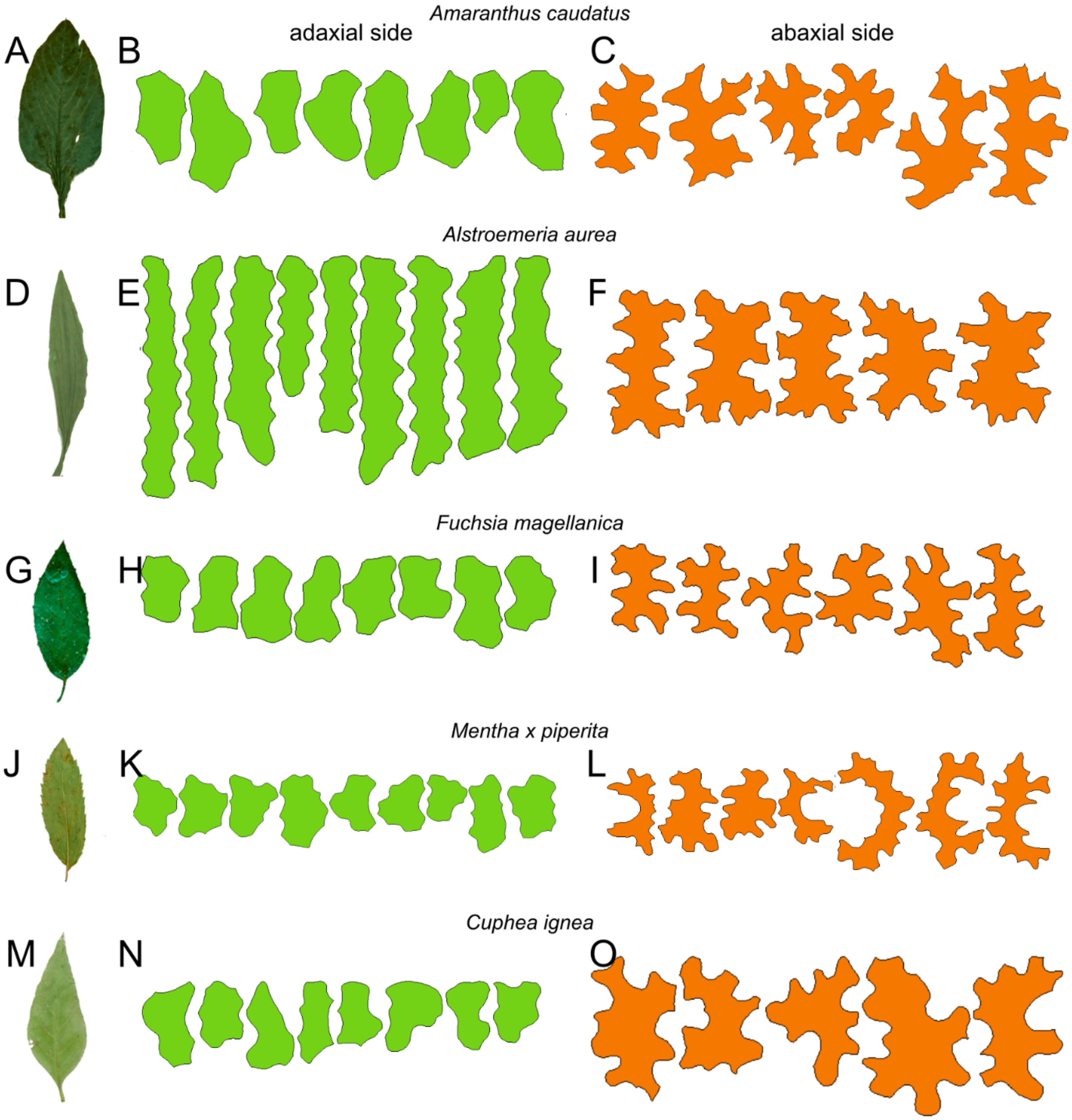


**Suppl. Figure 8. The lobeyness of pavement cells varies between adaxial and abaxial leaf surfaces.** (**A-O**) Leaf images (**A, D, G, J, M**) and corresponding pavement cell outlines representing the 95th percentile of lobeyness on the adaxial (cells colored in green) (**B, E, H, K, N**) and abaxial (cells colored in orange) (**C, F, I, L, O**) leaf surfaces. Each row represents a different species: love-lies-bleeding (*Amaranthus caudatus*) (**A-C**), Peruvian-lily (*Alstroemeria aurea*) (**D-F**), fuchsia (*Fuchsia magellanica*) (**G-I**), peppermint (*Mentha x piperita*) (**J-L**), and cigar flower (*Cuphea ignea*) (**M-O**). Scale bar for cell contours: 100 µm (**N, O**), 150 µm (**B, C, I, H, L**), 200 µm (**E, F, K**).

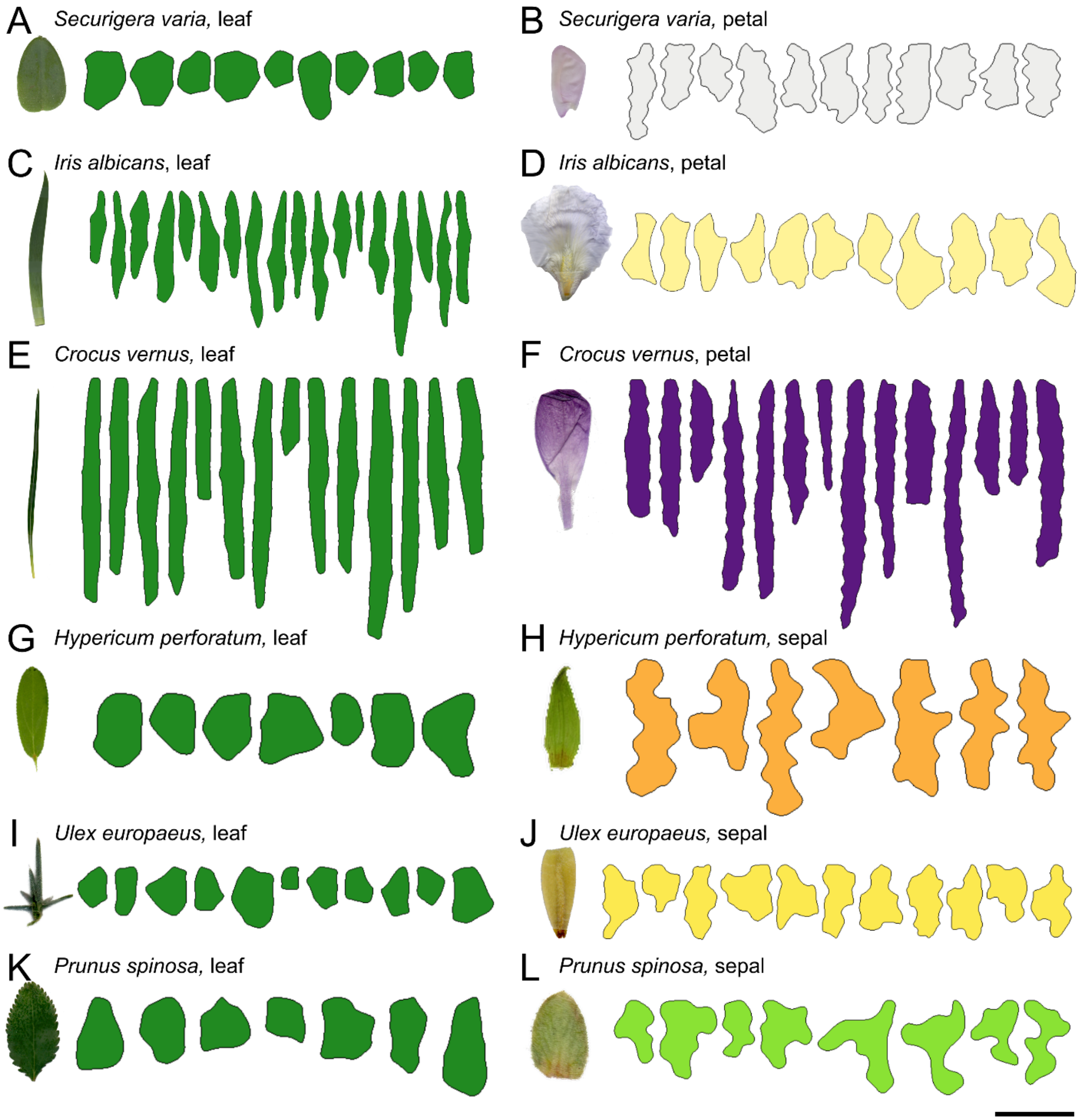


**Suppl. Figure 9. Organ-specific differences in pavement cell shape between leaves and floral structures.** (**A-L**) Leaf and corresponding non-lobed cell outlines (**A**, **C**, **E**, **G**, **I**, **K**) compared to lobed pavement cells of petals in crown-vetch (*Securigera varia*) (**B**), cemetery iris (*Iris albicans*) (**D**), spring crocus (*Crocus vernus*) (**F**), and sepals in St John's wort (*Hypericum perforatum*) (**H**), gorse (*Ulex europaeus*) (**J**) and blackthorn (*Prunus spinosa*) (**L**). Different colors indicate cells sampled from: adult leaves (dark green); petals and sepal contours are colored in accordance with the organ's appearance when possible. Scale bars for cell contours: 50 µm (**G**, **I**), 75 µm (**L**), 100 µm (**A-F**, **J**, **K**).

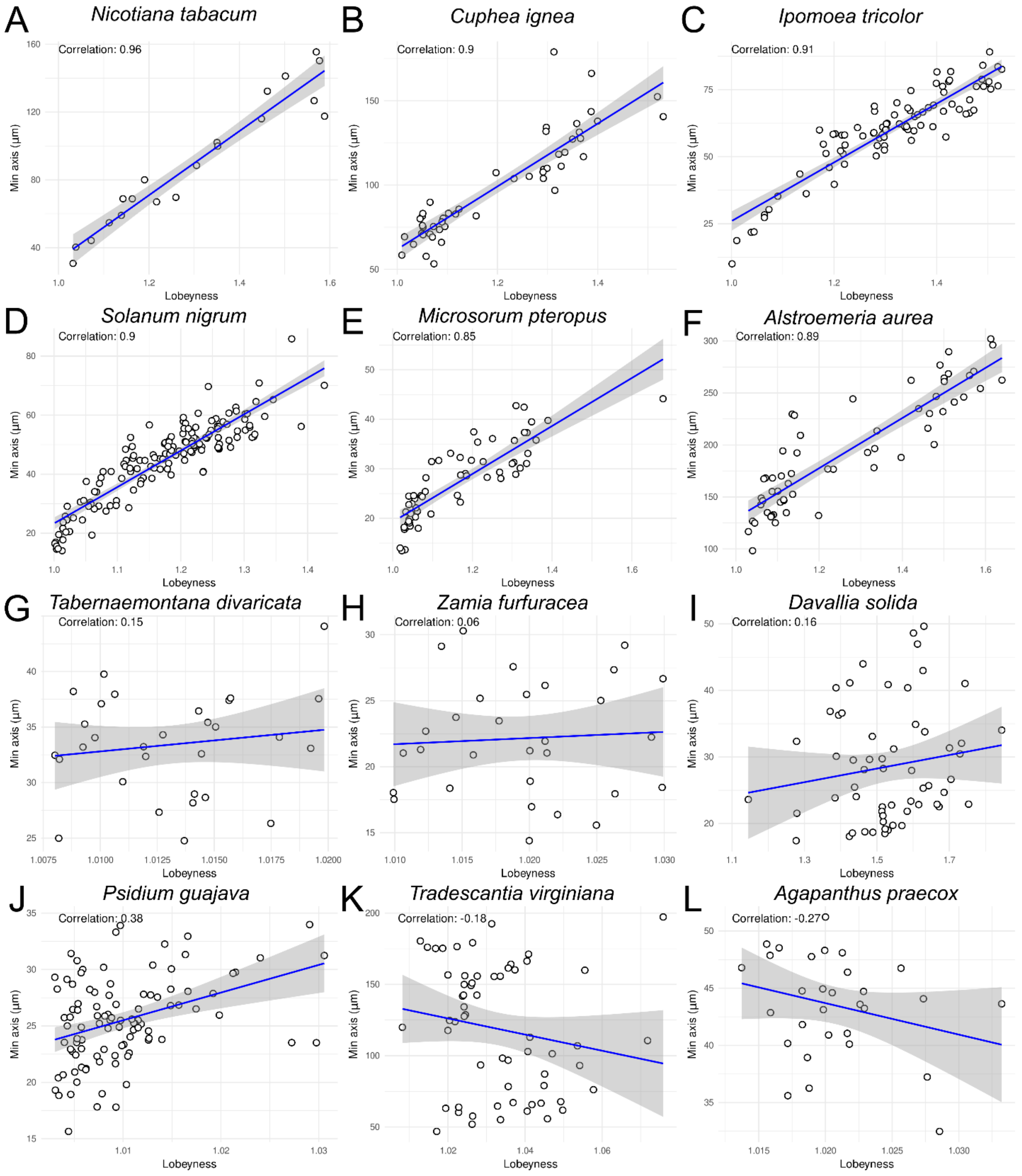


**Suppl. Figure 10. Correlation patterns between min-axis and lobeyness in pavement cells within individual species.** Each panel shows a scatter plot illustrating how the min-axis (the shortest cell dimension, y-axis) relates to lobeyness (x-axis) for a given species, revealing a spectrum of correlation strengths. (**A–F**) Examples of strong positive correlation in tobacco (*Nicotiana tabacum*) (**A**), cigar plant (*Cuphea ignea*) (**B**), morning glory (*Ipomoea tricolor*) (**C**), black nightshade (*Solanum nigrum*) (**D**), Java fern (*Microsorum pteropus*) (**E**), and Peruvian lily (*Alstroemeria aurea*) (**F**). (**G–L**) Weak or negative correlation in pinwheel flower (*Tabernaemontana divaricata*) (**G**), cardboard palm (*Zamia furfuracea*) (**H**), hare's foot fern (*Davallia solida*) (**I**), yellow guava (*Psidium guajava*) (**J**), Virginia spiderwort (*Tradescantia virginiana*) (**K**), and agapanthus (*Agapanthus praecox*) (**L**).

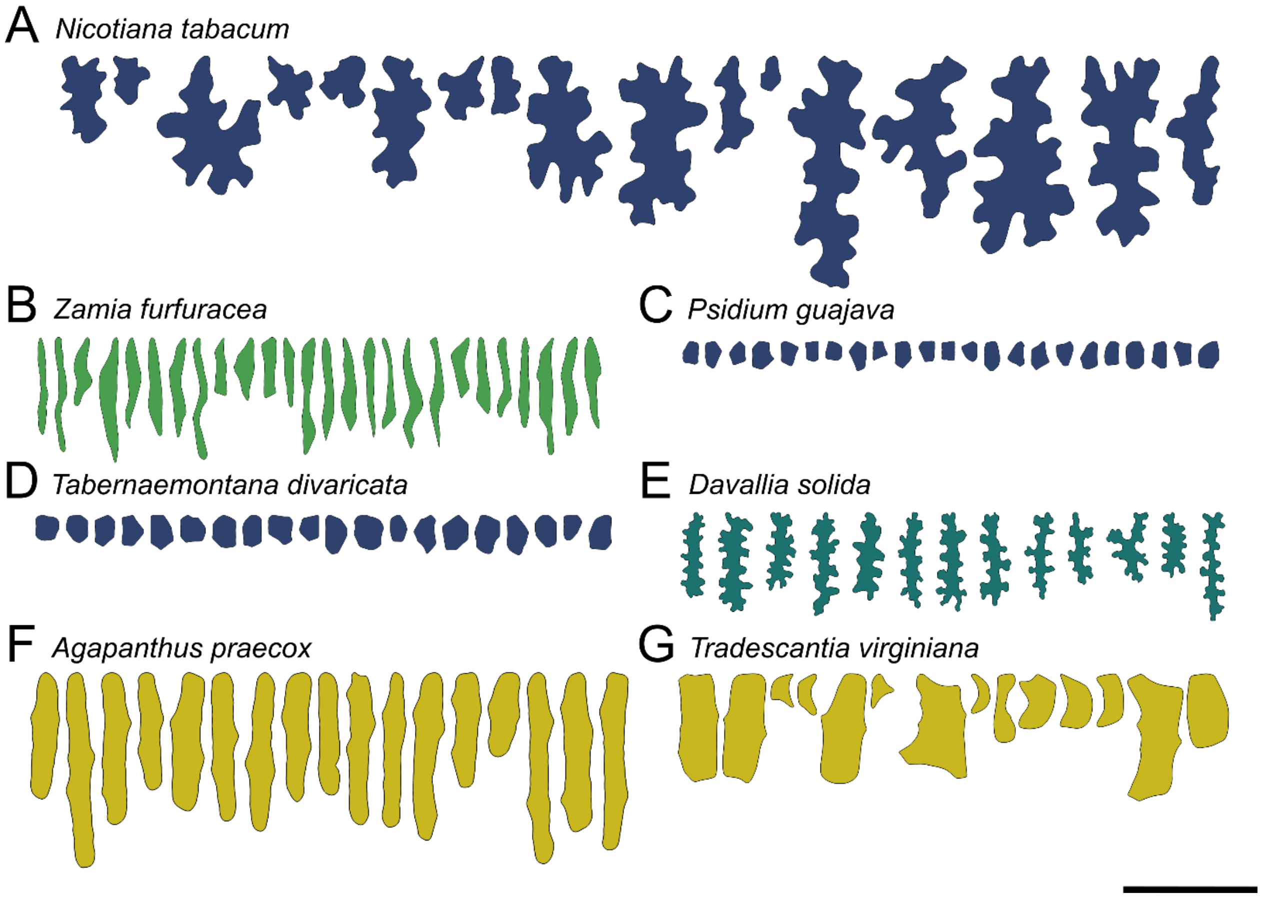


**Suppl. Figure 11. Visual examples of pavement cell contours demonstrating high, low, and negative correlations between lobeyness and min-axis.** (**A-G**) Cell contours of leaves color-coded by clade. (**A**) Pavement cells of tobacco (*Nicotiana tabacum*) that have a high correlation between min-axis and lobeyness (Corr = 0.97). (**B**) The cardboard palm cells (*Zamia furfuracea*) have a low correlation that is typical for long and thin cells (Corr = 0.24). (**C**, **D**) Epidermal cells in yellow guava (*Psidium guajava*) and pinwheel flower (*Tabernaemontana divaricata*) maintain small cells of a uniform size (Corr = 0.38 and Corr = 0.13, respectively). (**E**) Highly lobed cells in the hare's foot fern (*Davallia solida*) display little variation in lobeyness, which influences the correlation (Corr = 0.16). (**F**) In agapanthus (*Agapanthus praecox*), bumps next to junctions of neighboring cells affect thinner cells more than thicker ones, resulting in negative correlations (Corr = -0.21). (**G**) Negative correlations are also typical for pavement cells in the Virginia spiderwort (*Tradescantia virginiana*) with the much smaller stomatal lineage cells displaying irregular concave shapes (Corr = -0.2). Scale bars: (**A-D**, **F**) 200 µm, (**G**) 250 µm.

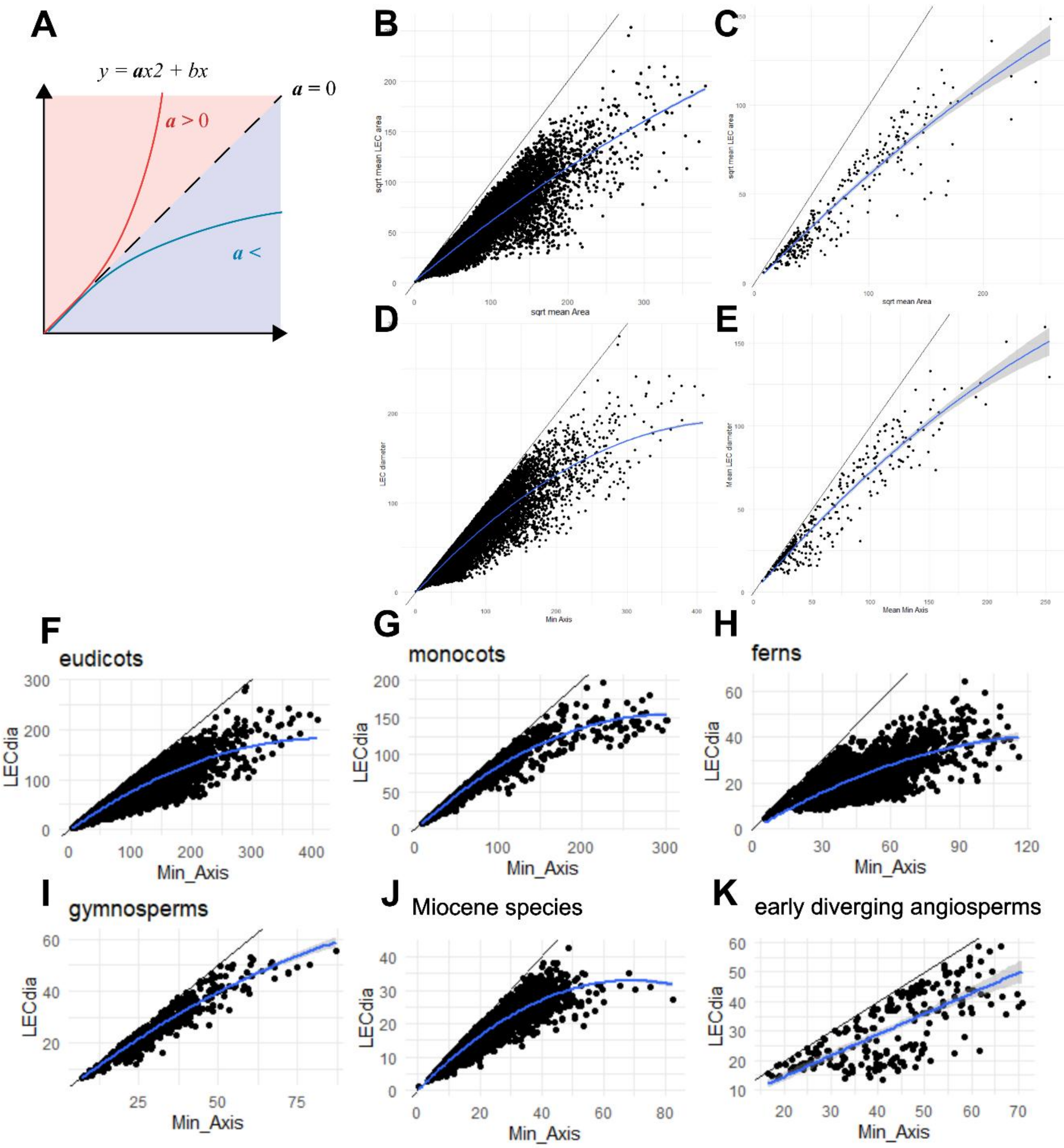


**Suppl. Figure 12. The analysis of LEC across different species suggests a correlation between cell size and lobeyness.** The black line in each plot indicates the theoretical limit where LEC area equals cell area (**B, C**), or LEC diameter equals min-axis length (**D-K**). The blue line indicates the best fit quadratic polynomial passing through the origin, grey shaded region indicates the 95% confidence interval. (**A**) The sign of the coefficient on the quadratic term indicates whether LEC tends to increase more quickly ($\alpha$>0) or slowly ($\alpha$<0) as cell size increases. (**B-C**) LEC area vs cell area for all cells (**B**) and species means (**C**). (**D-E**) LEC diameter vs min-axis length for all cells (**D**) and species means (**E**). (**F-K**) LEC diameter vs min-axis for all cells in the indicated clade. Negative α values were observed in 87% of all species (286 of 386 species, p < 2e-16<0 .05, exact binomial test), eudicots 87% (172/197 species, p < 2e-16<0 .05); monocots 78% (38/49 species, p = 1.4e-4<0 .05); ferns 97% (40/41 species, p = 3.8e-11<0 .05); gymnosperms 84% (16/19 species, p=4.4e-3<0 .05); Miocene species 100% (13/13 species, p=2.4e-4<0 .05); early diverging angiosperms 100% (7/7 species, p=0.01<0.05).

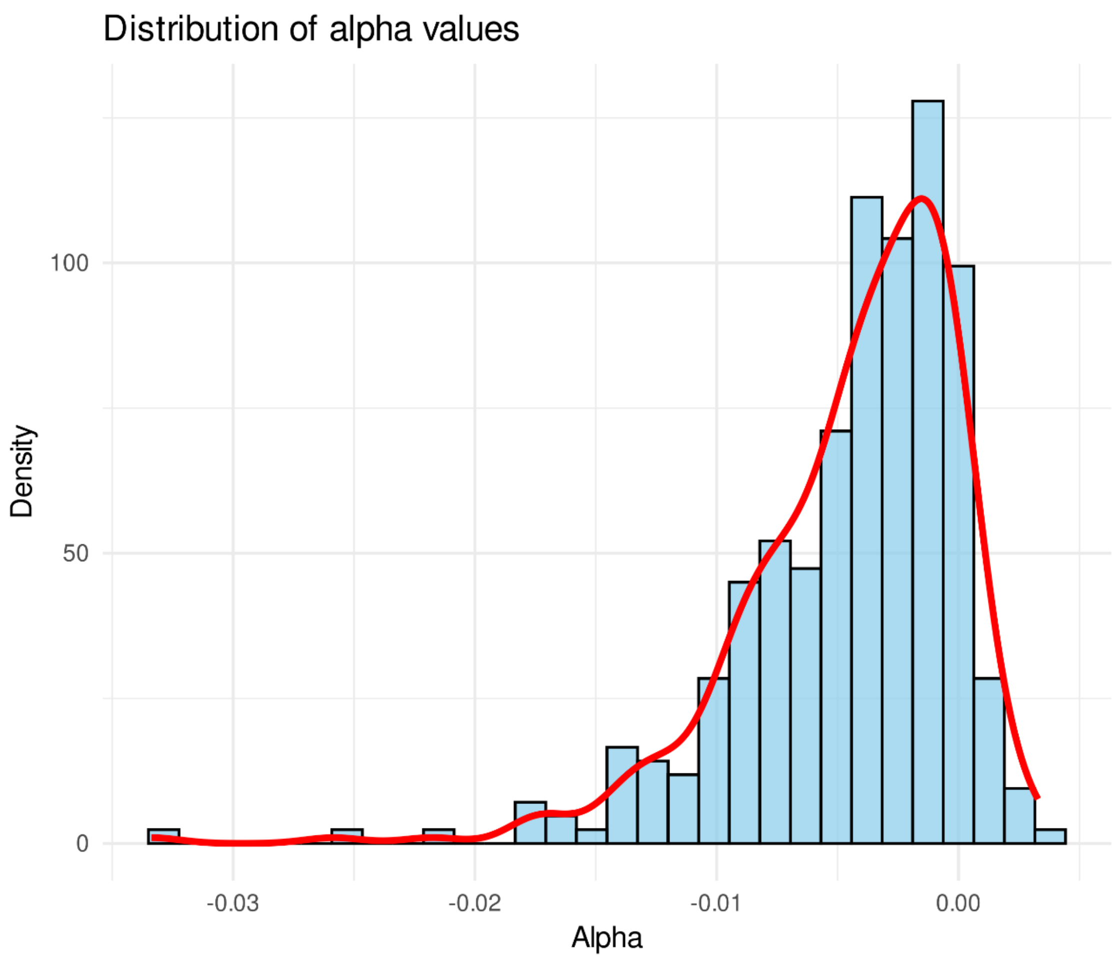


**Suppl. Figure 13. Distribution of alpha values across species.** The histogram shows the distribution of alpha values (bins = 30) computed from the quadratic fit of LEC versus min-axis. The red density curve represents smooth distribution. Most alpha values are negative, supporting the conclusion that LEC increases at a slower rate as the min-axis becomes larger. An exact sign-test was performed to assess the probability of observing 286/386 negative coefficients by chance, confirming that this pattern is highly unlikely to occur randomly.

### Material and Methods:

#### Plant material and growth conditions

The seeds were sterilized by immersing them in a solution of 70% EtOH (Sigma-Aldrich, Cat: 64-17-5) and TWEEN® 20 (Sigma-Aldrich, Cat: P9416) for 2 minutes. Subsequently, they were rinsed twice with 95% EtOH and left to air dry. The sterilized seeds were then placed in square Petri plates (Sigma-Aldrich, Cat: Z692344) filled halfway with a growth medium composed of ½ MS (Murashige and Skoog Basal Salt Mixture, Duchefa Biochemie, Cat: M5524), 1% sucrose (Fisher, Cat: 57-50-1), at pH 5.6, and 0.7% agar (Duchefa Biochemie, Cat: P1001). After a period of 72 hours of stratification at 4°C, the seeds were grown in a vertical position for 5 days at 22°C, with a 16-hour light cycle per day, in a controlled environment room. Subsequently, the 5-day-old seedlings were transferred to soil and cultivated for 4 weeks at 22°C, with a 16-hour light cycle per day. The screen of epidermal pavement cell outlines was conducted on *Arabidopsis thaliana* Col-0 ecotype and the following lines: *anisotropy1* (*any1*) [1], *constitutively active GTP-bound ROP2* (*CA-ROP2*) [2], and *constitutive triple response1* (*ctr1*) [3]. For the microtubule imaging, seeds of green fluorescent protein – α-TUBULIN6 (GFP-TUA6) line [4] were grown on a Petri plate for 12 days and stained with 0.1% PI prior imaging. For the cell wall elasticity and turgor pressure measurements, pAR169 (pATML1::mCitrine-RCI2A) line was used to visualize plasma membrane, and mCit-MBD (pPDF1::mCit-MBD) line was used for visualization of microtubules.

#### Confocal imaging of the cell outlines and image processing

The epidermal pavement cell outlines were imaged from the adaxial surfaces of 3-week-old *Arabidopsis thaliana* leaves. Small sections (5 $mm^2$) were sampled from the central part of the leaves and submerged in a staining solution of 0.1% Propidium Iodide (PI) (Sigma, Cat: 81845) dissolved in water for 20 minutes. After a brief rinse in water, the samples were mounted on a slide with a cover slip and imaged using a Leica STELLARIS upright laser confocal microscope equipped with a water immersion objective (25x/0.95). The excitation and emission wavelength windows used were 488 nm and 600-650 nm for PI and 488 nm and 495-545 nm for GFP, respectively. The images were acquired with 1024x1024 resolution and z-stack slices with 0.5-1 μm in the z-direction. At least 10 different plants and leaves from the adaxial sides were collected at the same developmental stage for each genotype. The confocal images were processed in MorphoGraphX using *Gaussian Blur Stack* and *Brighten Darken* filters to enhance the clarity of cell outlines. The confocal signal was then projected onto a mesh, reconstructing the three-dimensional structure of the sample (2.5D). Cell segmentation was performed using seeding and *Watershed Segmentation* processes. To better capture the cell size aspect relevant to stress, we introduced the min-axis parameter. Unlike cell area, the min-axis accounts for the smallest dimension of the cell by measuring the width of the minimum bounding rectangle

(**Figure 3A**). This approach allows for a more accurate representation of cell size in the context of lobing, as elongated cells with large areas but small LECs are not expected to form lobes. The min-axis parameter distinguishes between long, thin cells (low min-axis values) and cells that are large in both dimensions (high min-axis values). Lobeyness, calculated as the ratio of a cell's perimeter to its convex hull perimeter (the reciprocal of convexity), was employed to assess lobing. Heatmaps for cell parameters including lobeyness, min-axis, area, and LEC were generated using MorphoDynamX. Correlation analysis between contour parameters was conducted using the MDXtoR plugin.

### Sampling of different plant species

The analysis of epidermal cell outlines from different species (**see Suppl. Table 1**) was conducted using specimens collected from commonly grown species in Cologne (Germany) and Norwich (United Kingdom). To obtain the cell outlines, we employed the imprinting method with 3% Low Melting Point (LMP) agarose (ROTH, Germany, Cat: 39346-81-1) dissolved in dH2O. The agarose solution was briefly microwaved until boiling and then allowed to cool to room temperature without solidifying. On a microscope slide, a droplet of the lukewarm agarose was carefully dispensed, small fragments (1 $cm^2$) were excised from the central region of the plant organs using a razor blade and then gently transferred onto the lukewarm agarose droplets and allowed to solidify. The plant sections were carefully removed from the slide while keeping the agarose intact, thus preserving the shape of the epidermal cells. The slides with the agarose imprints were subsequently stored at 4°C until they were imaged. Imprints were obtained from both the adaxial and abaxial sides of all collected organs, excluding green fruit, stamen, stigma, and bract. The identification of plant species was conducted using references such as Simpson [5] and Harris and Harris [6]. Refer to Sapala, et al. [7] and Vőfély, et al. [8] for the methods used for pre-existing contours.

### Phase-contrast microscopy of epidermal imprints

The transparent epidermal imprints, with a thickness of 5-10 µm, were visualized using either a Zeiss Axio Imager M2 microscope equipped with an Axiocam 512 camera, or a Nikon Microphot-SA microscope connected to a Leica camera. Air Plan-NEOFLUAR objectives of either x20 or x40 were utilized, depending on the cell size of the sample being imaged. To ensure representative coverage, each image typically contained approximately 50 cells, adjusted based on the specific sample. The phase-contrast mode was employed for imaging on both microscopes. The images were acquired using ZEN 2.3 software for the Zeiss Axio Imager M2 microscope, utilizing the tailing mode to stitch multiple images (5x5), or LAS software for the Nikon Microphot-SA microscope.

### Image acquisition for Miocene fossil plant species

The Miocene plant fossils analyzed in this study were obtained from a dataset published by Reichgelt *et al.* (2020). These fossils were preserved in turbidite deposits within the Foulden Maar diatomite core. To prepare the fossilized leaves for analysis, they were treated with hydrogen peroxide and tetra-sodium pyrophosphate salt crystals. This treatment effectively removed mesophyll cell debris, ensuring clearer observation of the leaf structures. The cleaned fossil leaves were then stained with Crystal Violet and mounted on glass slides using glycerin jelly. High-resolution images of the leaves were captured at 100× magnification using TSView 7.1.1.2 microscope imaging software on a Nikon Optiphot microscope. Species identification of the fossil specimens was based on paleobotanical studies and comparison with known species from the Foulden Maar surface exposures.

### Image processing for multispecies pavement cell outline analysis

To improve the visualization of cell outlines, GNU Image Manipulation Program (GIMP) was used to enhance saturation, contrast, and sharpness. Each image contained a maximum of 50 cells, and only cells with clear outlines were manually traced. Fiji software was used to calculate the distance in pixels along the scale length, which was used to calibrate the scale bars in MorphoGraphX [9]. To aid in cell segmentation, various image filters, including *Invert*, *Gaussian Blur Stack*, and *Brighten Darken*, were applied. The surface mesh was extracted using the *Marching Cubes Surface* algorithm with a cube size of 5 µm. The cell outlines from the phase-contrast microscopy images were projected onto the mesh, and cell segmentation was achieved by applying the *Watershed Segmentation process*. To further refine the cell contours, the *Smooth Mesh* process was applied, resulting in a smoother and more accurate representation of the cells. The cell contours obtained from the segmented mesh were then analyzed and quantified using MorphoDynamX. A subset of the most representative cell contours was selected for further analysis, ensuring a focus on the cells that were most relevant to the research objectives. Heatmaps for lobeyness, min-axis, area, and LEC were generated using the *Contour process*. Correlation analysis between two contour parameters was performed using the MDXtoR plugin.

### Dose-dependent drug treatment

To investigate the effects of oryzalin or latrunculin B on *Arabidopsis thaliana* cotyledon pavement cells, plates containing 50 mL of MS medium (0.7% agar, without sugar and vitamins) were prepared. The drugs, oryzalin (Sigma-Aldrich, Cat: 36182) or latrunculin B (Sigma-Aldrich, Cat: L5288), were added to the plates at final concentrations of 0.5, 1, 5 and 8 µM. As a control, DMSO (Dimethyl sulfoxide, Sigma, Cat: D8418) was added to the growth medium at a volume equivalent to that used for the drug treatments. After 6 days, the cotyledons were collected and mounted on microscope slides with cover slips. Images of the cotyledon pavement cells were captured using Leica STELLARIS upright laser confocal microscope equipped with a water immersion objective (25x/0.95). The captured images were then processed to segment the cells and quantify their contours.

### Growth quantification for epidermis of juvenile maize leaves

Maize (*Zea mays* cv. 'Polonez' and 'Cosmo') caryopses were kept submerged in water for 6 h, germinated on blotting paper for 18 h in light at room temperature, transferred to plastic containers filled with moist soil, and grown in glasshouse (temperature 19-21 °C, additional illumination to obtain 16 h day). After 4-7 days, a longitudinal strip of coleoptile or coleoptile and the first leaf sheath was gently removed to expose the base of the first or second juvenile leaf, respectively. The sequential replica method [10,11] was used to obtain silicon moulds, made of Take 1 Advanced Impression Material (Light Body Wash, Kerr Corp., Romulus, USA) from the abaxial epidermis of the exposed leaf surface. Two replicas were taken from each leaf at 24 h interval. After replica taking the exposed leaf portion was protected from drying by food wrap. Epoxy resin replicas (casts made from Devcon 2 Ton Clear epoxy) were obtained from the silicon moulds, sputter-coated and observed using scanning electron microscopy (Hitachi UHR FE-SEM SU8010). Pairs of stereoimages were taken from each region of interest and used for the stereoscopic reconstruction of the leaf surface [12]. Growth was analysed for 2 juvenile leaves of cv. 'Polonez' and 3 juvenile leaves of cv. 'Cosmo'. For each leaf, 3-12 patches of epidermis were identified in the two consecutive replicas, located at an increasing distance from the intercalary meristem. Because maize pavement cells become strongly elongated, in each patch, 3-9 groups (nearly square-shaped in the first replica), which comprised 6-12 adjacent cells, were used to compute areal growth rates and Principal Growth Directions [13].

### Contour loading and pre-processing

Contours from Vőfély, et al. [8] (www.doi.org/10.5061/dryad.g4q6pv3) were loaded into MorphoDynamX (www.MorphoDynamX.org) using a custom process written for this purpose (Mesh/Contours/Vofely Load Contours). Each cell was given a unique label, and then labelings (one to many groupings) were assigned based on clade, species, and species-side. Contours were scaled so that units are in microns and smoothed with a single pass position average of their 1-neighborhoods to reduce noise. A small number of contours were encountered with topological errors, such as cells in multiple pieces or with vertices out of order causing self-intersection. The very few cells in multiple pieces were removed, and a process was written to select vertices out of order. These specific vertices were smoothed again, and the process repeated, which removed any remaining problems.

### Contour processes

Several new processes were added to MorphoDynamX to aid in contour processing. There processes are as follows:

*Arrange Contours* - Arrange contours in a grid for visualization. This process also sorts them by a specified heatmap, or the current if nothing specified, for example, lobeyness.

*Create Contours* - Create contours from a 2.5D surface mesh. This simplifies cells made of multiple faces into a single planar face and rotates them into the XY plane.

*Create LEC* - Create a cell complex with the LEC visualized. The cell label and heat are preserved.

*Load Contours* - Loads contours from a directory as text files. Contour files are specified as a list of 2D positions, one per line, with the units assumed to be microns.

*Load Contours Labeled* - Loads contours from a directory structure root/clade/species/organ/sample.

*Rotate Min-Axis* - Rotates contours so that the min-axis is in the X direction.

*Save Contours Labeled* - Saves contours to a directory structure root/clade/species/organ/sample.

*Select Bad Contours* - Selects points on contours where there are topological problems.

*Select by Percentile* - Selects the specified percentile cell for the specified labeling.

*Trim Contours* - Remove selected points from contours.

### Min-axis calculation

For a 2D contour, the min-axis is the axis along which the contour has minimum width. To accelerate computations, we use the convex hull of the contour, which substantially reduces the complexity of the contour without affecting its width with respect to a given axis. To find the min-axis we densely sample axis orientations between 0 and 2π, and project the convex hull onto each axis to determine the contour's width. Once the min-axis has been identified, we set its length to be equal to the contour's width along this axis.

### Mechanical model

The simulation model of puzzle shape emergence was adapted from Sapala, et al. [7]. The model is a fully damped mass-spring mechanical model written in C++ using the MorphoDynamX modeling framework (www.MorphoDynamX.org). In this model, cells are represented by point masses (vertices) linked together by linear springs. Based on the cell shape, additional intracellular connections (springs) are placed representing the orientation of microtubules that guide the deposition of cellulose within the cell. Importantly, these connections are restricted to locations where geometric constraints, such as connection angle and curvature, permit their formation. The model can thus be used to predict the orientation and distribution of microtubules based on the outlines of cell walls obtained through microscopic imaging.

Growth is simulated by fixing the positions of the outermost vertices, moving them slightly outward at each simulation step before finding the positions of the inner vertices at the equilibrium between the forces on springs and the intracellular connection constraints. The rate at which the outer vertices are moved was varied to simulate distinct growth patterns. For uniform growth, the outer vertices move apart at a constant rate, possibly different in the horizontal and vertical directions, throughout the simulation. For nonuniform growth, the rate of movement of the outer vertices changes as the simulation runs.

The main changes to the model for this work were to add functions to allow for changing growth over time in the X and Y directions. These functions use the format of the "funcedit" function editing tool supplied with the vlab modeling environment (http://algorithmicbotany.org/virtual_laboratory/). This tool creates a function file (.func) that contains spline points to define the function. The function file can be edited interactively with funcedit or with a text editor. Step functions were created to simulate a period of full growth followed by a period of zero growth (StepDown.func) and vice-versa (StepUp.func). Growth functions for the maize simulation were fit to growth data (ZeaGrowthX.func, ZeaGrowthY.func). A description of the model parameters, starting templates and growth processes can be found in the "description.txt" file contained in the model.

### Data availability

Experimental data, simulation models, additional MorphoDynamX processes and the MorphoDynamX software are available online on Dryad. See the ReadMe.txt file for a description of the archived contents.

**The following plant organ scans are modified from:**

*Amaranthus caudatus*: NEU000004163. Neuchâtel Herbarium.
*Alstroemeria aurea*: BR0000025021905. Meise Botanic Garden Herbarium.
*Fuchsia magellanica:* 304262. Field Museum of Natural History.
*Mentha x piperita:* 00058444P. Oxford University Herbarium.
*Cuphea ignea:* 289029. Oxford University Herbarium.
