## Supplemental Table 1 for "Growth history leaves a geometric trace in puzzle cells"

**Suppl. Table 1.** Species used for the analyses classified by clade.

| early diverging angiosperms | eudicots | ferns | gymnosperms | Miocene species | monocots |
|---|---|---|---|---|---|
| Annona montana | Acanthus hungaricus | Actiniopteris semiflabellata | Araucaria | Araliaceae-Pseudopanax | Acorus gramineus |
| Chloranthus | Acanthus spinosus | Adiantum hispidulum | Araucaria bidwillii | Atherospermatacea-Laurelia otagoensis | Agapanthus praecox |
| Drimys winteri | Acer platanoides | Angiopteris | Bowenia serrulata | Elaeocarpaceae-Cunoniaceae | Agave sisalana |
| Illicium anisatum | Achillea millefolium | Angiopteris evecta | Chamaecyparis thyoides | Lauraceae-Cryptocarya maarensis | Albuca bracteata |
| Laurus nobilis | Aconitum carmichaelii | Asplenium adiantoides | Cycas revoluta | Lauraceae-Cryptocarya sp | Aloe |
| Magnolia denudata | Adromischus | Asplenium lancifolium | Encephalartos transvenosus | Lauraceae-Cryptocarya taieriensis | Alpinia zerumbet |
| Magnolia lotungensis | Alliaria petiolata | Asplenium vieillardii | Ginkgo biloba | Lauraceae-Endiandra | Alstroemeria aurea |
| Peperomia | Alternathea dentata | Bolbitis | Gnetum montanum | Lauraceae-Litsea calicarioides | Amaryllis belladonna |
| | Amaranthus caudatus | Bolbitis lonchophora | Lepidozamia peroffskyana | Meliaceae-Dysoxylum | Amorphophallus titanum |
| | Amaranthus hybridus | Ceratopteris thalictroides | Microcycas calocoma | Myrtaceae | Avena strigosa |
| | Anchusa officinalis | Cyrtomium falcatum | Picea abies | Primulaceae-Myrsine | Calibanus hookeri |
| | Anemone canadensis | Davallia fejeensis | Picea glauca | Ripogonaceae-Ripogonum | Carex morrowii |
| | Anthemis arvensis | Davallia solida | Podocarpus macrophyllus | Unknown-Morphotype O | Carludovica palmata |
| | Ardisia crispa | Davallia subsolida | Taxodium distichum | | Clivia miniata |
| | Astragalus falcatus | Dicksonia antarctica | Taxus canadensis | | Crocus sativus |
| | Bellis perennis | Diplazium plantaginifolium | Tsuga canadensis | | Crocus vernus |
| | Bergenia purpurascens | Doodia media | Widdringtonia nodiflora | | Cymbidium |
| | Berkheya purpurea | Elaphoglossum | Zamia furfuracea | | Danae racemosa |
| | Berkheya radula | Goniopteris | Zamia pumila | | Danthonia californica |
| | Beta trigyna | Helminthostachys zeylanica | Zamia skinneri | | Dioscorea bulbifera |
| | Beta vulgaris | Lecanopteris sinuosa | | | Dioscorea mexicana |
| | Borago officinalis | Lygodium japonicum | | | Eichhornia crassipes |
| | Bougainvillea | Lygodium microphyllum | | | Eriachne |
| | Brugmansia suaveolens | Marattia attenuata | | | Eustrephus latifolius |
| | Bupleurum fruticosum | Microsorum pteropus | | | Freycinetia |
| | Bursera schlechtendalii | Nephrolepis | | | Haworthia retusa |
| | Callicarpa japonica | Niphidium crassifolium | | | Hemerocallis fulva |
| | Calotropis gigantea | Osmunda banksiifolia | | | Hordeum vulgare |
| | Calystegia sepium | Pellaea paradoxa | | | Hydrocleys nymphoides |
| | Campanula fenestrellata | Pellaea viridis | | | Iris |
| | Campanula poscharskyana | Phanerophlebia | | | Iris albicans |
| | Capsella bursa-pastoris | Phlebodium aureum | | | Iris japonica |
| | Capsicum annuum | Platycerium andinum | | | Kniphofia caulescens |
| | Cardamine flexuosa | Platycerium elephantotis | | | Leucojum aestivum |
| | Carica papaya | Polypodium | | | Oryza sativa |
| | Cassia spectabilis | Pteris ensiformis | | | Pandanus tectorius |
| | Cataranthus roseus | Rumohra adiantiformis | | | Rohdea japonica |
| | Catha edulis | Sphaeropteris cooperi | | | Ruscus aculeatus |
| | Catharanthus roseus | Stenochlaena palustris | | | Ruscus hypoglossum |
| | Ceiba pentandra | Tectaria | | | Sabal minor |
| | Centaurea cineraria | Tectaria pseudosinuata | | | Setaria italica |
| | Centaurea cyanus | Todea barbara | | | Smilax bona-nox |
| | Centranthus ruber | | | | Sorghum bicolor |
| | Cephalaria flava | | | | Sternbergia lutea |
| | Ceratostigma plumbaginoides | | | | Stichoneuron caudatum |
| | Ceratostigma willmottianum | | | | Tradescantia virginiana |
| | Cerinthe minor | | | | Tricyrtis hirta |
| | Ceropegia sandersonii | | | | Triraphis |
| | Chenopodium bonus-henricus | | | | Yucca gloriosa |

| | |
|---|---|
| Choisya ternata | Zea mays |
| Cinnamomum camphora | |
| Cissus quadrangularis | |
| Cissus tiliacea | |
| Citrus limon | |
| Clerodendrum thomsoniae | |
| Cordia nitida | |
| Crassula | |
| Crataegus monogyna | |
| Cuphea ignea | |
| Dahlia pinnata | |
| Dillenia indica | |
| Dodonaea viscosa | |
| Dorycnium rectum | |
| Epiphyllum | |
| Ercilla spicata | |
| Eryngium agavifolium | |
| Eryngium bourgatii | |
| Erysimum scoparium | |
| Erythrina standleyana | |
| Euphorbia flanaganii | |
| Euphorbia mellifera | |
| Euphorbia pulcherrima | |
| Fagraea berteroana | |
| Forsythia suspensa | |
| Fraxinus excelsior | |
| Fuchsia magellanica | |
| Fuchsia triphylla | |
| Fuschia Mrs Popple | |
| Galanthus nivalis | |
| Galium odoratum | |
| Galium rubioides | |
| Geranium pusillum | |
| Geum triflorum | |
| Globularia punctata | |
| Globularia trichosantha | |
| Grevillea flexuosa | |
| Guaiacum officinale | |
| Haloragis erecta | |
| Hedera nepalensis | |
| Heimia myrtifolia | |
| Helleborus orientalis | |
| Helminthotheca echioides | |
| Hiptage benghalensis | |
| Homalocladium platycladum | |
| Hypericum patulum | |
| Hypericum perforatum | |
| Ilex aquifolium | |

Ilex paraguariensis

Impatiens balsamina

Impatiens repens

Ipomea tricolor

Jacquemontia tamnifolia

Jasione heldreichii

Jasminum fruticans

Jasminum humile

Justicia guttata

Lactuca sativa

Lamium orvala

Lamium purpureum

Leucanthemum vulgare

Linaria vulgaris

Lithocarpus henryi

Lonicera quinquelocularis

Macleania insignis

Malva sylvestris

Medicago sativa

Mentha x piperita

Mespilus germanica

Mimosa pudica

Montinia caryophyllacea

Murraya koenigii

Mussaenda glabra

Myrrhis odorata

Nemophila menziesii

Nicotiana tabacum

Nyctanthes arbor-tritis

Oenothera glazioviana

Oenothera stricta

Othonna capensis

Oxalis latifolia

Oxalis triangularis

Oxalis valdiviensis

Oxyria digyna

Paeonia tenuifolia

Papaver rhoeas

Passiflora

Passiflora edulis

Pastinaca sativa

Pelargonium carnosum

Penstemon

Persicaria polystachya

Persicaria weyrichii

Physalis origanifolia

Plantago afra

Poliothyrsis sinensis

Polygonum affine

Populus tremula x tremuloides

Potentilla reptans

Prunus spinosa

Psephellus simplicicaulis

Psidium guajava

Rauvolfia verticillata

Rhus potaninii

Ribes sanguineum

Rosa x damascena

Rubia tinctorum

Rumex acetosella

Rumex scutatus

Sambucus nigra

Saxifraga canaliculata

Saxifraga hostii

Scabiosa olgae

Scorzonera hispanica

Scrophularia canina

Scrophularia heterophylla

Securigera varia

Senna didymobotrya

Silene latifolia

Sisymbrium austriacum

Solanum nigrum

Stachys macrantha

Strobilanthes wallichii

Succisella inflexa

Symphytum caucasicum

Symphytum grandiflorum

Symphytum orientale

Syringa

Syringa vulgaris

Tabernaemontana divaricata

Taraxacum officinale

Tecoma stans

Thunbergia mysorensis

Tilia cordata

Torrenia asiatica

Trifolium pannonicum

Ulex europaeus

Valeriana phu

Verbena bonariensis

Veronica gentianoides

Veronica missurica

Veronica persica

Veronica petraea

Veronica rakaiensis

Viola odorata
